# MacSyFinder v2: Improved modelling and search engine to identify molecular systems in genomes

**DOI:** 10.1101/2022.09.02.506364

**Authors:** Bertrand Néron, Rémi Denise, Charles Coluzzi, Marie Touchon, Eduardo P.C. Rocha, Sophie S. Abby

## Abstract

Complex cellular functions are usually encoded by a set of genes in one or a few organized genetic loci in microbial genomes. Macromolecular System Finder (MacSyFinder) is a program that uses these properties to model and then annotate cellular functions in microbial genomes. This is done by integrating the identification of each individual gene at the level of the molecular system. We hereby present a major release of MacSyFinder (version 2) coded in Python 3. The code was improved and rationalized to facilitate future maintainability. Several new features were added to allow more flexible modelling of the systems. We introduce a more intuitive and comprehensive search engine to identify all the best candidate systems and sub-optimal ones that respect the models’ constraints. We also introduce the novel *macsydata* companion tool that enables the easy installation and broad distribution of the models developed for MacSyFinder (macsy-models) from GitHub repositories. Finally, we have updated and improved MacSyFinder popular models: TXSScan to identify protein secretion systems, TFFscan to identify type IV filaments, CONJscan to identify conjugative systems, and CasFinder to identify CRISPR associated proteins. MacSyFinder and the updated models are available at: https://github.com/gem-pasteur/macsyfinder and https://github.com/macsy-models.

## INTRODUCTION

Microbial machineries and pathways (hereafter called “systems”) can be very complex and involve many proteins. In the genomes of Bacteria and Archaea, the proteins constituting these systems are often encoded in a highly organized way, involving one or a few operons with functionally related genes. For example, loci encoding the peptides of a protein complex or the enzymes of a metabolic pathway have specific genetic organizations that tend to be remarkably conserved (Dandekar et al., 1998; Teichmann & Babu, 2002). Neighbouring operons in genomes are also often functionally related (Huynen et al., 2000). This means that gene co-localization can be used to infer gene functions and improve homology inference, *e*.*g*., when sequence similarity is low. Co-localization also facilitates the distinction between functionally diverging homologs (Abby & Rocha, 2012). The hypothesis is that the member of the gene family that co-localizes with the rest of the system’s genes is the one involved in the functioning of this system. In addition, many cellular processes require the involvement of a coherent ensemble of proteins. In such cases, the genetic potential for a function can only be identified when the repertoire of genes is analyzed at the system-level. For example, a minimum set of proteins (and thus of encoding genes) is necessary for the functioning of a protein secretion system.

In 2014, we published the “Macromolecular System Finder” (MacSyFinder v1) program for the functional annotation of cellular machineries and metabolic pathways in microbial genomes (Abby et al., 2014). It makes a system-level annotation that takes advantage of the typical functional organization of microbial genomes (gene co-localization) and the requirement of a core set of proteins to perform the function (quorum). Such concepts have already been successfully applied in other annotation tools, such as KEGG mapper (quorum) (Kanehisa & Sato, 2019), Antismash (quorum and co-localization) (Blin et al., 2021) or Pathways Tool (Karp et al., 2020) for accurate annotations of enzymes participating in metabolic pathways. Yet these tools were created to make specific metabolic annotations, and were not designed for the user to develop their own annotation tools for any type of macromolecular system. MacSyFinder consists of a generic modelling framework and a search engine to screen genomes for candidate systems. The modelling framework enables the user to define models for the systems of interest, including the corresponding genes’ identity, category, and genetic organization. MacSyFinder v1 has three categories of genes: mandatory, accessory, and forbidden. Parameters of gene co-localization describe the genomic architecture of the system at the level of each gene or of the entire system. Each gene corresponds to one HMM (hidden Markov model) protein profile to enable sequence similarity search with HMMER (Eddy, 2011), and different genes (thus protein profiles) can be defined as exchangeable if they have the same role in the system. The search engine screens a database of genomes for potential systems using HMM profile searches and the clustering of co-localized hits along the genome that match the systems’ model.

MacSyFinder has been used with success to annotate a variety of microbial machineries and pathways, including protein secretion systems (Abby et al., 2016), CRISPR-Cas systems (Abby et al., 2014; Couvin et al., 2018) and other prokaryotic defence systems (Tesson et al., 2022), capsular loci (Rendueles et al., 2017), DNA conjugation systems (Cury et al., 2020), the butyrate production pathway (Sharp & Foster, 2022), methanogenic and methylotrophic metabolisms (Adam et al., 2019; Chibani et al., 2022), cell division machineries (Pende et al., 2021) and outer membrane protein clusters (Taib et al., 2020). It has enabled wide-scale genomic analyses of biologically relevant systems and was integrated into the popular MicroScope genome annotation pipeline and in the reference CRISPRCasFinder program (Couvin et al., 2018; Vallenet et al., 2020).

In spite of its successful applications, MacSyFinder v1 has several limitations. In terms of software engineering, it is coded in the now obsolete Python v2.7, lacks tools to improve its future development and maintenance, and some parts of the program are not efficient. In terms of modelling, it cannot use gene-specific criteria to filter the HMMER hits when annotating the genes of a system. Furthermore, it lacks a way to annotate genes of interest that are neutral concerning the systems’ assessment. More importantly, the greedy search engine is not optimal and has complex, sometimes counter-intuitive behaviours.

We hereby present a major release of MacSyFinder, MacSyFinder version 2 (v2) coded in Python 3 (>= 3.7). In addition, we have updated and improved the most popular MacSyFinder models to the new version to make them readily usable.

## MATERIALS AND METHODS

### Input & Output files

MacSyFinder v2, like v1, takes as input files the models of the systems to search and a multi-protein FASTA file (Figure 1). The user defines the search mode that corresponds to the nature of the input FASTA file. When all the proteins encoded by a replicon are ordered in the file following the genes’ order, one can use the most powerful search mode – “ordered_replicon” – to include both the analysis of the genetic content (quorum) and of the genetic organization. Otherwise, the search mode “unordered” is used, and only the genetic content is assessed.

**Figure 1.**
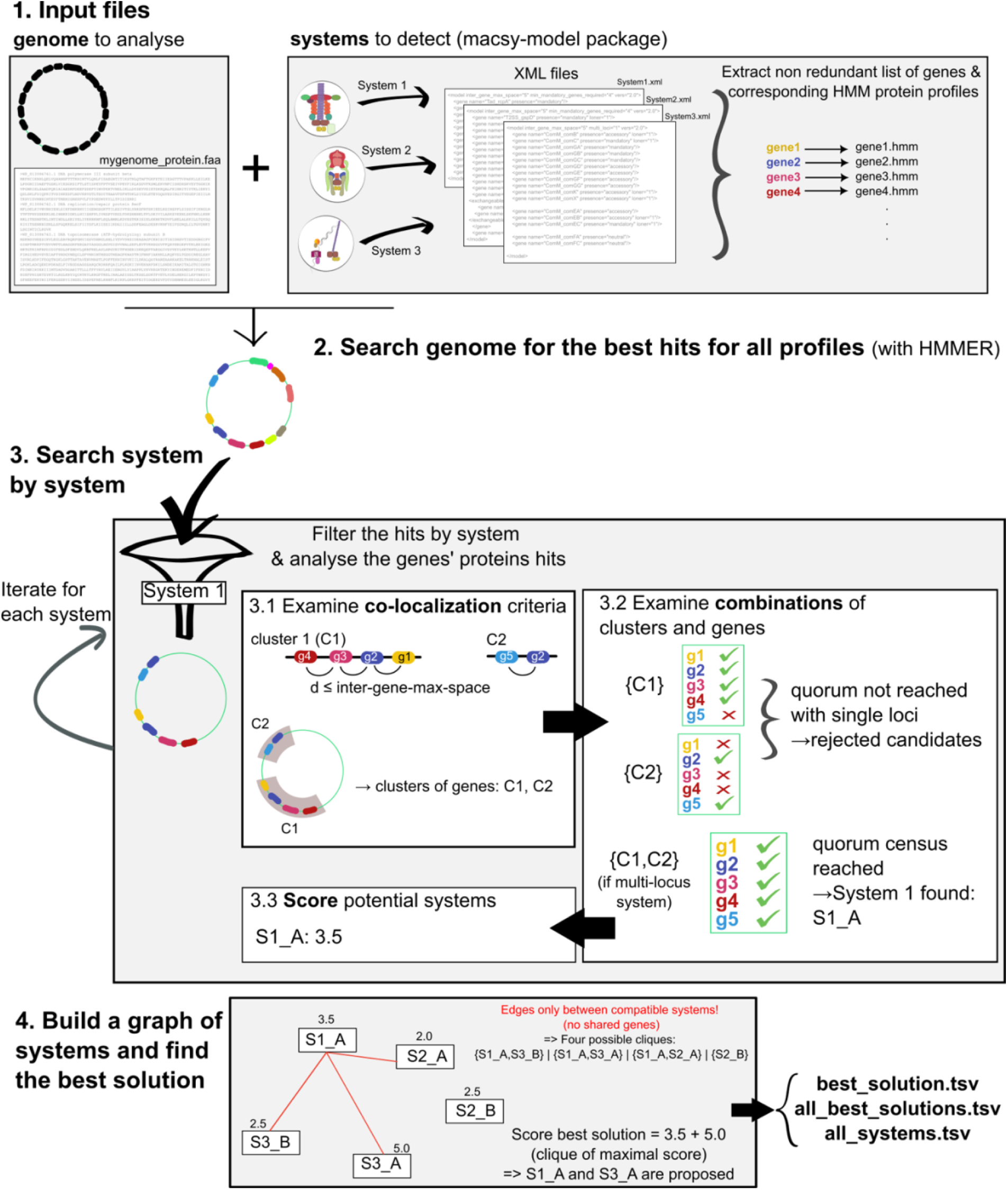
Overview of the major steps of MacSyFinder v2. The user gives as input the genome(s) to analyse under the form of a multi-protein FASTA file (order respecting that of the genes on genome if possible) and a macsy-model package with the systems to detect. Then the search engine establishes the non-redundant list of corresponding genes. (2) The genes are then searched with HMMER (hmmsearch using GA scores when available) using the corresponding HMM protein profiles. In the absence of GA score, the proteins with the best hits are filtered by i-evalue and profile coverage (if no GA score was available). (3) A system-by-system search is then performed. The hits corresponding to a first system are selected (“g1” as hit for gene 1), and clusters of genes are formed by gathering the hits respecting the maximal inter-gene-max-space (3.1). Genes allowed to be “out-of-clusters” are also collected (loners and multi-systems). Then the possible combinations of clusters and “out-of-clusters” genes are computed, and the program tests if they respect the quorum for the system (3.2). Finally, all candidate systems are scored (3.3). Step (3) is re-iterated for each system to be detected. Once all systems have been examined, the best solution is searched using a graph-based approach. Step (4) A graph connecting all compatible candidate systems (i.e., systems with no shared genes) is built, each node having the score of the corresponding system. The best solution is defined as the set of compatible systems obtaining the highest cumulative score. This corresponds to the clique of maximal score. Diverse output files are provided to the user, including one with the composition of (one of) the best solution, a file with all equivalent best solutions (if several reach the highest score), and one with all eligible candidate systems whether they are part of the best solution or not. Drawings of systems at Step (1) are derived from (Denise et al., 2019).

Of note, recommendations on how to use MacSyFinder on incomplete or fragmented genomes are included in the “How To” section of the User guide. In a nutshell and depending on the level of assembly and completeness of the genome, we recommend to run MacSyFinder with the “ordered_replicon” mode, which can be complemented by the results of an “unordered” run. Results using the “ordered replicon” option on draft genomes have to be considered with care.

A third option is to use the “gembase” mode. This requires that multiple ordered genomes are provided in a single FASTA file using headers with a pre-determined naming convention (see documentation). The program Panacota (Perrin & Rocha, 2021) can provide such a database. A Nextflow workflow “*parallel_macsyfinder*” is provided at the MacSyFinder GitHub repository to enable the analysis of multiple genomes in parallel based on a “gembase” file (see the User guide https://macsyfinder.readthedocs.io/en/latest/user_guide/big_data.html).

The significant modifications in terms of input and output in v2 concern the organization of the input systems’ models (“macsy-model” packages, see below) and the output files. The latter were adapted to reflect the new MacSyFinder search engine results. In addition, various easy-to-parse text tabulated files are now proposed as output, including the raw and filtered results of the proteins’ similarity search with HMMER, the gene-wise description of the possible systems, the systems constituting the best solutions, and the gene-wise description of rejected candidates. For more details, one can consult MacSyFinder’s comprehensive documentation, including the User Guide, the Modeller Guide, and the Developer Guide, created with Sphinx and available at: https://macsyfinder.readthedocs.io/. Two datasets showing command line examples and expected input and output files are provided on the Figshare platform: https://doi.org/10.6084/m9.figshare.21581280 and https://doi.org/10.6084/m9.figshare.21716426.v1.

### Formalizing macsy-model packages

MacSyFinder v1 required two directories containing one or several systems’ models (“definitions” folder) and the corresponding HMM profiles (“profiles” folder). These files were passed to the command line as mandatory parameters and were distributed as standalone archives. Unfortunately, these had the inconvenience of being poorly versionable or traceable. To improve the reproducibility of analyses with MacSyFinder v2, we increased the traceability of the models and facilitated their retrieval and installation by formalizing a package structure that we call “macsy-model” package (see Fig. S1).

A macsy-model package must have two directories: “definitions” and “profiles”. The “definitions” directory contains all model definitions written in the MacSyFinder-specific XML grammar (one file per model definition). This directory can include several sub-directories and levels (Fig. S1). The “profiles” directory contains all HMM protein profiles included in the definitions. In addition, a new file, “*metadata*.*yml”*, was introduced to store necessary metadata such as the package name, version, description, citation, distribution license, and the contacts of its author(s)/maintainer(s). Some facultative but recommended files can be added: LICENSE/copying, Contributing, README.md, and model_conf.xml. The README.md file should explain how to use the models and can be displayed using the command *macsydata help* (see below). The file *model_conf*.*xml* allows the modeller to set package-specific configurations such as score configuration options (see paragraph on scoring) or criteria to filter the hits when searching the genes’ proteins (profile coverage threshold, usage of GA scores with HMMER, etc.). The user can easily supersede these recommended values using the command line and configuration files.

### Grammar update for the modelling framework

The models of MacSyFinder are written using a dedicated XML grammar with a hierarchy that fits the hierarchical nature of the biological systems to model: systems’ models are made of gene components (Fig. 3, Supplementary Table 1). The two main objects in the hierarchy of a system’s model are thus the “model” (replacing the “system” keyword in v1) at the top level and the “gene” at the lower level. In addition, a feature “vers” was added at the model level to indicate the version of the grammar: “vers=2.0” matches MacSyFinder v2.

We simplified the XML grammar to ensure better readability and easier maintenance of the models (Fig. 3, Supplementary Table 1). Relative to the first version, some keywords were removed or merged into novel ones. This is the case of the keywords “homologs” and “analogs” that were replaced by the new keyword “exchangeables” to indicate that some genes can be “exchanged” by others (*i*.*e*., fill the same role in systems). The gene attribute “exchangeable” was thus removed as not needed anymore. The “system_ref” keyword was also removed.

When designing a system’s model or investigating the distribution of genes within genomic occurrences of a system, one may want to annotate genes that are not important to the identification/discrimination of the system but provide information on accessory functions. To achieve this, we introduced a new type of gene called “neutral”. It adds to the pre-existing types - mandatory, accessory, and forbidden - with the key difference that it is not used to score the systems or to assess their quorum (minimal number of genes required in a system). Neutral genes are identified using HMM protein profiles and placed into clusters like the other genes. Hence, even if they do not contribute to the scoring of systems, they can “connect” or “extend” clusters of mandatory and accessory genes.

Details, examples, and a tutorial concerning the XML grammar v2 are available in MacSyFinder’s documentation, which now includes a novel section called the “Modeller Guide” to explain how to build novel models of systems. This is also the focus of our recent book chapter that covers all aspects of the modelling process (Abby et al., 2023). We hereby provide some of the most popular models translated with improvements for MacSyFinder v2 (Table 1). They are readily usable and installable through the *macsydata* program (see Results section).

**Table 1.** Overview of MacSyFinder v2 macsy-model packages available at the “MacSy models” organization https://github.com/macsy-models

| Model repository | Version tag | Original references | Systems detected | No models | No profiles | Remark |
| --- | --- | --- | --- | --- | --- | --- |
| CasFinder | 3.1.0 | (Abby et al., 2014; Couvin et al., 2018) | CRISPR-Cas systems<br>(Cas clusters detection, annotation, and classification) | 44 | 535 | This new version:<br>- provides the possibility to detect more subtypes than the previous ones<br>- greatly improves the detection of tandem systems<br>- improves the detection of decayed and atypical systems<br>- can now be ran at once for the three levels of classification<br>- GA scores added to HMM profiles |
| CONJscan | 2.0.1 | (Cury et al., 2017) | Conjugative, mobilizable and decayed conjugative systems | 34 | 124 | This new version:<br>- allows the detection of decayed conjugative systems<br>- provides models adapted to the detection in chromosomes or plasmids<br>- contains tailored thresholds for each type of MPF<br>- GA scores added to HMM profiles |
| TFFscan | 1.0.0 | (Denise et al., 2019) | Systems members of the type IV filament super- family | 7 | 169 | As in the original paper, but with GA scores added to HMM profiles |
| TXSScan | 1.0.1 | (Abby et al., 2016) | Protein secretion systems and related appendages | 22 | 205 | As in the original paper, but with GA scores added to HMM profiles |
| TXSScan | 1.1.1 | (Abby et al., 2016; Denise et al., 2019) | Protein secretion systems and related appendages, including members of the type IV filament super-family | 26 | 341 | Merger of TFF-SF v1.0 and TXSScan v1.0:<br>- TFF-SF models from Denise et al. replace older model versions from TXSScan version 1.0 (Abby et al. ) for the T2SS, Tad and T4aP systems. Models added: ComM, T4bP and Archaeal_T4P.<br>- Hierarchy of models by domain of life, and then by membrane type to allow to search only the relevant models. |

### Enabling gene-wise filtering by setting up GA scores

The HMMER search for the systems’ genes can now use the “GA” (Gathering) bit scores of the corresponding HMM protein profiles. This score can be defined for each HMM profile to set a score threshold for the inclusion of the hit in the HMMER search results. The GA score is usually used as a minimal score limit for protein family inclusion (*e*.*g*. by PFAM). Using the GA score allows employing gene-wise criteria for protein hit filtering instead of having the same criteria for all genes (as in v1 of MacSyFinder). If a GA score is present in the HMM profile file, the system calls HMMER using the option “--cut_ga”, which supersedes the i-evalue (for “independent e-value”, a stringent type of e-value as computed by HMMER) and profile coverage default values otherwise used in the absence of GA scores. It is possible to deactivate the GA scores using the new option “--no-cut-ga” (False by default). The rules for the filtering of the protein hits can be specified (by decreasing order of priority): in the command line, in the model configuration file of the system (“model_conf.xml”), or using the HMM protein profile GA scores (or the “i-evalue” and “profile coverage” if GA scores not provided).

Many of the HMM protein profiles used in MacSyFinder models already include GA thresholds because they were retrieved from PFAM or TIGRFam, which systematically use them (Sonnhammer et al., 1997; Haft et al., 2003). Yet, some other profiles lacked GA thresholds. To remediate this limitation, we modified these profiles to include threshold GA scores. We did this for CasFinder, TXSScan, CONJScan, and TFFscan profiles (see Table 1). To this end, we annotated with the corresponding models the completely assembled genomes of 21105 bacterial and archaeal strains retrieved from the non-redundant NCBI RefSeq database (as of March, 2021, see Dataset S1). We analysed the distribution of the scores for the hits for the different genes found in the detected systems, and attributed as GA score the minimal score observed to the corresponding profile.

In total, 1000 HMM profiles from the four packages are available with GA scores. They now offer the possibility for gene-wise filtering with HMMER, ensuring optimal usage of the 104 macsy-models hereby updated for MacSyFinder v2.

### Sharing and handling macsy-models with the *macsydata* command and the “MacSy Models” organization

The novel tool “*macsydata*” was created to make macsy-model packages easily traceable, versionable, shareable, and automatically installable. It was designed to be as light as possible for the modellers and was inspired by the packaging workflows found in some Linux distributions such as Gentoo (https://www.gentoo.org). In addition, the *macsydata* API was inspired by *pip*, which is familiar to most Python users. *macsydata* implements common sub-commands such as *search, install, upgrade, uninstall*, and *help* (see Table 2). We also implemented some specific useful sub-commands for MacSyFinder, such as *cite* to display how to cite the macsy-model package, *definition* to show a set of models’ definitions in XML format, or *init* to initialize a template for a new macsy-model package.

**Table 2.** The *macsydata* companion tool to handle macsy-model packages

| *macsydata* command | Description | Examples |
| --- | --- | --- |
| *available* | Lists all macsy-model packages available at the default or specified organization | *macsydata available* |
| *list* | Lists installed packages | *macsydata list* |
| *init* | Initializes a new macsy-model package with a template of the appropriate file architecture | *macsydata init* |
| *check* | Allows to check the sanity and consistency of a macsy-model package before diffusion | *macsydata check* |
| *install / uninstall* | Automatically retrieves and installs (uninstall) the designated package | *macsydata install* TFF-SF |
| *cite* | Displays the citation information stored in the metadata.yml file of the package | *macsydata cite* TFF-SF |
| *definition* | Displays the definition(s) (XML file) of the specified model(s). It can be a directory containing several models to display. | *macsydata definition* TXSS T1SS<br>*macsydata definition* TXSS/archaeal<br>*macsydata definition* –models-dir my-models System1 |
| *search* | Searches for the models based on their names, or based on string searches in the models' description | *macsydata search* TXSS<br>*macsydata search* -s Secretion |
| *macsydata*<br>*<subcommand> --help* | Lists the help message for the specified sub-command | *macsydata search --help* |

The “MacSy Models” Github organization was designed to serve as an umbrella organization to host any macsy-model package. It allows the modeller to efficiently distribute its packages to all MacSyFinder v2 users. Firstly, one must create a *git* repository and develop a macsy-model package *e*.*g*., following the Modeller guide or the recommendations in (Abby et al., 2023). Secondly, the quality of the package (package structure, model definitions syntax, the coherence between definitions and profiles) can be checked using the *macsydata check* command on the directory containing the entire file architecture of a macsy-model package (Fig. S1). Finally, when everything is up to standards, the modeller has just to tag the repository and push it under the Github organization “MacSy Models”. This action allows the model package to be visible from the *macsydata search* tool and thus findable and accessible for remote installation to all MacSyFinder users. *macsydata* uses the Github Rest API to search and download the packages. Of note, *macsydata* can also install macsy-model packages from a tarball archive, given it respects the above-described file architecture.

### The macsyprofile companion tool

The novel tool “*macsyprofile*” of the MacSyFinder suite allows filtering and extracting HMMER hits with settings different from those used during the run. This allows retrieving relevant hits not initially included in predicted systems, *e*.*g*., to understand why they were “missed”. This could be particularly useful to assist the design of the profiles and the systems’ models or to search for atypical versions of the systems (see details in the online documentation).

### Comparison of MacSyFinder v1 and v2

MacSyFinder version 1.0.5 and MacSyfinder v2.0.0 were run on the dataset of complete bacterial and archaeal genomes described above (RefSeq March 2021) to detect the TXSS and related systems. The models of TXSScan v1.1.1 were used with MacSyFinder v2 while the models published in (Denise et al., 2019) for the TFF-SF and (Abby et al., 2016) for the other secretion systems were used with MacSyFinder v1. The total number of systems detected in the dataset were compared between the two MacSyFinder versions for each type of system. The same was done to compare the median system wholeness (proportion of the model-listed genes detected) for each system.

### Testing the performance of MacsyFinder v2

We evaluated the performance of MacSyfinder by measuring the running time and the RAM used. We measured the overall time of the run and the time spent in several parts of the software: the genes identification by HMMER (hmmsearch) and the resolution of the best solution. To assess the used RAM, we prefixed the *macsyfinder* command line by the “/usr/bin/time -v” utility and extracted the “Maximum resident set size” value. We ran the analysis on a sub-set of the above-described complete genomes dataset, consisting of one genome per bacterial species. This corresponded to 6092 genomes (6455 chromosomes, see Dataset S1). To analyze the behavior of the resolution of the best solution, we ran *macsyfinder* with these genomes as input and using the “ordered_replicon” mode with three macsy-model sets: TXSScan/bacteria, CasFinder and CONJscan/chromosome. We also compared the novel algorithm based on graphs’ usage (using NetworkX) to an implementation based on a Mixed-Integer Linear Solver (Python-MIP) (Supplementary Figures 4 & 5). To assess the time spent in each part of the program, we ran *macsyfinder* with the TXSScan/bacteria models in “gembase” mode on datasets gathering increasing numbers of genomes (1, 10, 20, 50, 100, 200, 500) selected from the 6092 genomes set. All tests were performed on the following setup: Python version 3.11, linux kernel version 5-15.80, AMD Ryzen 7 3700X 8-Core Processor (16 threads) for CPU, and 64GB DDR4 for RAM.

### Code implementation, dependencies, and availability

The code was ported to Python 3 (>=3.7). Many unit and functional tests were implemented to reach a coverage of the code of 97%. In terms of dependencies, the program requires the HMMER suite (>=3.1b2) for the search of proteins. It also uses several well-established and stable Python libraries to facilitate models’ packaging (*pyyaml, packaging*), deal with output files (*colorlog, pandas*), and search for the best solution (*NetworkX*, see below).

The code and macsy-model packages are available on Github under the GPL v3 license: https://github.com/gem-pasteur/macsyfinder and https://github.com/macsy-models. In addition, a pypi package, a conda package and a Docker container were created to enable the easy deployment of the MacSyFinder suite.

## THE MACSYFINDER V2 SEARCH ENGINE

An overview of the new search engine is provided in Figure 1. The first steps of MacSyFinder v2 search engine remain mostly unchanged relative to v1. First, it uses HMMER to search for occurrences of the non-redundant genes listed in the models in the input genome(s) using the corresponding HMM protein profiles. The best hits are assigned to the corresponding genes and are filtered by profile coverage (>50% by default) and i-evalue (<0.001 by default) when no GA score is available for the profiles.

### System-wise creation of candidate systems

In version 2, the systems are searched one by one: the identified proteins have their hits filtered by type of system, and clusters of the corresponding genes are built from genes respecting the co-localization criteria (“inter-gene-max-space” parameter) (Fig. 1). Candidate systems are built using the clusters of genes and the genes authorized to be outside of clusters (“loner” genes). For “single-locus” systems, combinations of individual clusters with loner genes not yet represented in the cluster are examined as candidate systems. For systems allowed to be encoded by multiple loci, all possible combinations of identified clusters and loner genes (not found in clusters) are assessed as candidate systems. The eligible systems are the candidate systems that respect the systems’ model in terms of the minimal quorum criteria for all genes and the mandatory ones. The other systems are rejected for now and kept aside. In the case where “multi-system” genes are part of the systems’ model, the list of the corresponding genes will be collected from the eligible systems and combinatorically added to the set of rejected candidates to be assessed for the formation of new eligible systems.

### Introducing a scoring scheme for candidate systems

Candidate systems that respect the quorum and co-localization conditions imposed by a system’s model are designated as eligible systems and are assigned a score (Fig. 2A). The core of the system score is the sum of three terms: the sum of the scores of the *n* Clusters, the sum of the scores of the *o* out-of-cluster genes (loner or multi-system, see below), plus a penalty *P*_*system*_ for the redundancy within the system.

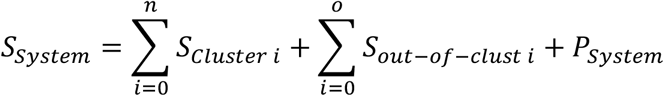

**Figure 2.**
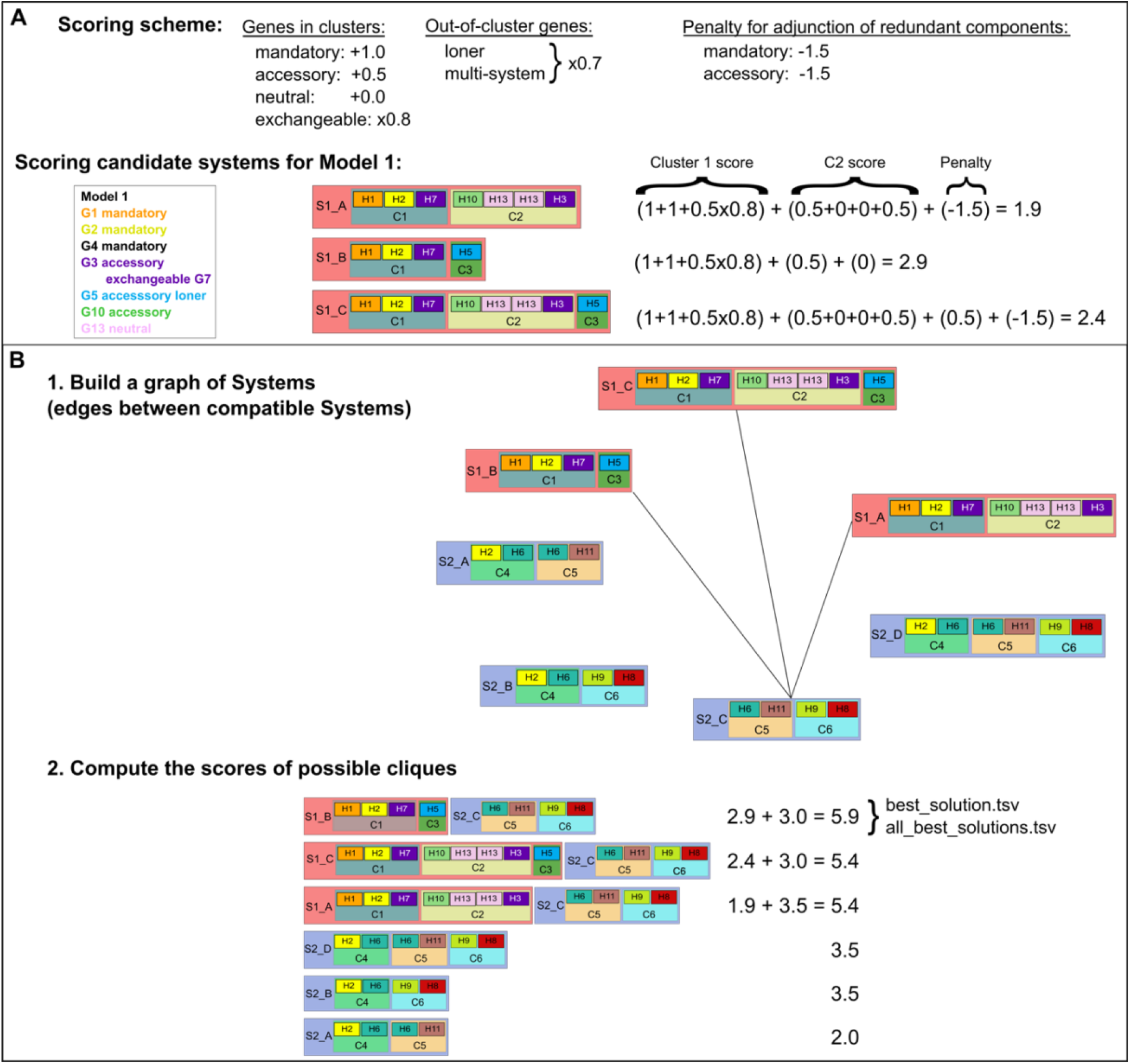
Scoring scheme and combinatorial search of MacSyFinder v2 search engine. **A**. The scoring scheme is summarized, and then illustrated by an example for a hypothetical Model1. “H1” stands for a hit for gene 1 “G1” in the genome. **B**. Step (1). The graph of candidate systems is drawn by connecting all compatible systems, *i*.*e*. those with non-overlapping hits (unless authorized by the multi-model or multi-system feature). Step (2). The clique of maximal cumulated score (best solution) is searched, with the score being defined as the sum of the systems’ scores that are part of the clique. The results are stored in the files “best_solution.tsv” and “all_best_solutions.tsv”, and all candidate systems are stored in “all_systems.tsv”.

Multiple occurrences of a gene within the same cluster are counted as a single occurrence of the gene. The score of each cluster is a function of the number of mandatory (*m)* and accessory genes (*a*), and of the number of exchangeable mandatory (*x*_*m*_) and exchangeable accessory (*x*_*a*_) genes it contains. These values are weighted to give more importance to mandatory genes: *w*_*mandatory*_ *= 1, w*_*accessory*_ *= 0*.*5*, and *w*_*neutral*_ *= 0* (Fig. 2A). Moreover, to give more value to the originally listed gene than to the listed “exchangeables”, a factor *f*_*exchang*_ *= 0*.*8* is applied to the scores when an exchangeable gene fulfils the function. The score *S*_*Cluster*_ is then given by:

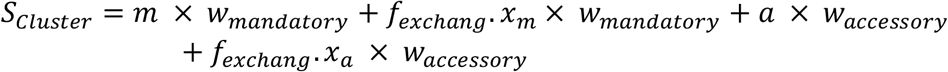

The score of the genes found outside of a system’s cluster is computed like the score of the genes found in clusters *“s*_*c*_*”* (see above), except that a factor *f*_*out-of-clust*_ *=* 0.7 is applied. Here, *s*_*c*_ can represent any of the gene-specific parts of the *S*_*Cluster*_ sum presented above, depending on the mandatory, accessory, and/or exchangeable status of the out-of-cluster gene:

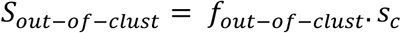

A gene is deemed redundant only if found in more than one cluster. The penalty part of a score thus penalizes candidate systems with *r* redundant mandatory or accessory genes, where *r* thus corresponds to the number of clusters with the gene minus one. We define *P*_*System*_ (*p*_*redundant*_ = −1.5 by default):

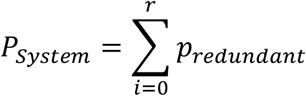

The default values of the different score parts are indicative and allow MacSyFinder to give priority to mandatory over accessory components, and to favour the main listed genes over the ones listed as “exchangeables”. Here the relative order of the values are more important than the absolute ones, and the users can fully parameterize the weights, factors, and penalties. The modeller of a system can also ship, with a macsy-model package, its recommended values for the weights of the scoring system using the optional “*model_conf*.*xml*” file.

### Combinatorial exploration of solutions

Once all models are searched and their occurrences are assigned scores (see above), the program performs a combinatorial examination of the possible sets of compatible systems is performed (Fig. 1 and 2B). Two systems are deemed compatible if they are made of distinct gene sets. Thus, unless specified using the “multi_system” or “multi_model” features, a gene cannot be involved in several systems. A MacSyFinder search solution is defined as a set of compatible systems. The search for the best solution corresponds to the well-known weighted maximum clique search problem (Brandes & Erlebach, 2005). The program builds a graph where each node represents a system with its associated score (as a weight), and where only compatible systems are connected with an edge. The goal is to identify a set of systems that are all compatible with each other, meaning that they are all inter-connected in a sub-graph. A sub-graph where all nodes are inter-connected is the definition of a “clique”. The best solution is thus the clique harboring the highest cumulated systems’ score, the score of a solution being the sum of the systems’ scores composing it. The “find_cliques” function proposed in the NetworkX Python library is used to find the set of maximal cliques that correspond to the best possible solutions in terms of cumulated nodes’ weights of the cliques (Hagberg et al., 2008). This may result in several solutions with the maximal score, in which case they are all provided to the user (file “all_best_solutions.tsv”). The best solution, or one among the best, is given in the dedicated output file “best_solution.tsv”.

## RESULTS & DISCUSSION

### I/ Grammar changes and macsy-model file architecture enable better, simpler, and more intuitive systems’ modelling and sharing

Version 1 of MacSyFinder lacked a dedicated file architecture to share MacSyFinder’s systems’ models. We now define a structured file architecture for the novel “macsy-model packages” (see Materials and Methods and Fig. S1). In particular, there now may be several levels of sub-directories for the “definitions” folder. This enables running *macsyfinder* with only a pre-defined subset of models and establishing a hierarchy of models in a biologically relevant manner. The introduction of this file architecture thus satisfies two main objectives: it allows the file architecture of the macsy-model packages to reflect the biological specificities of the systems while enabling automated handling of the macsy-model packages for easier distribution and installation via the *macsydata* tool. Several popular MacSyFinder models from v1 were ported to MacSyFinder v2 grammar and file architecture. They are now available at the “MacSy Models” Github organization for automated installation with MacSyFinder v2 using the *macsydata* tool (discussed in detail below, see also Materials and Methods, Table 1 and Table 2). The creation of the “MacSy Models” organization enables the macsy-model packages to be versioned for better reproducibility. This organization also constitutes the first step towards unifying a MacSyFinder modeller community.

To illustrate the interest of the novel file architecture, we created a new version of “TXSScan” (v1.1.1) that gathers the models for the type IV filament super-family (“TFF-SF”) and for the protein secretion systems (former “TXSScan”, v1.0.1) (Abby et al., 2016; Denise et al., 2019). These models were also ported to the grammar of MacSyFinder v2. The systems in “TXSScan v1.1” represent a coherent set of bacterial appendages dedicated to motility and secretion that are evolutionarily related (see (Denise et al., 2020) for a review). We organized the models into sub-directories with respect to relevant biological criteria (Fig. S1). The models’ sub-directories were split by domains of life (Archaea versus Bacteria) and then by membrane type (monoderm Bacteria versus diderm Bacteria). This new architecture enables the search for all models at once, or only the domain-specific ones or those specific to a given bacterial membrane type. This allows more targeted and less costly searches for biological systems. Of note, the XML grammar simplifications introduced in v2 enabled the production of much more compact, readable, and simple models (Fig. 3). For example, the definition of the type III secretion system (T3SS) now consists of 20 lines for 15 listed genes, whereas the v1 version counted 52 lines. v2 versions of the popular TXSScan and TFF-SF models are also available as originally published (v1.0.0 versions of TXSScan and TFFscan respectively, Table 1) (Abby et al., 2016; Denise et al., 2019). Yet the new search engine has a different behaviour than v1; it will produce different results in some cases (detailed in sections below).

**Figure 3.**
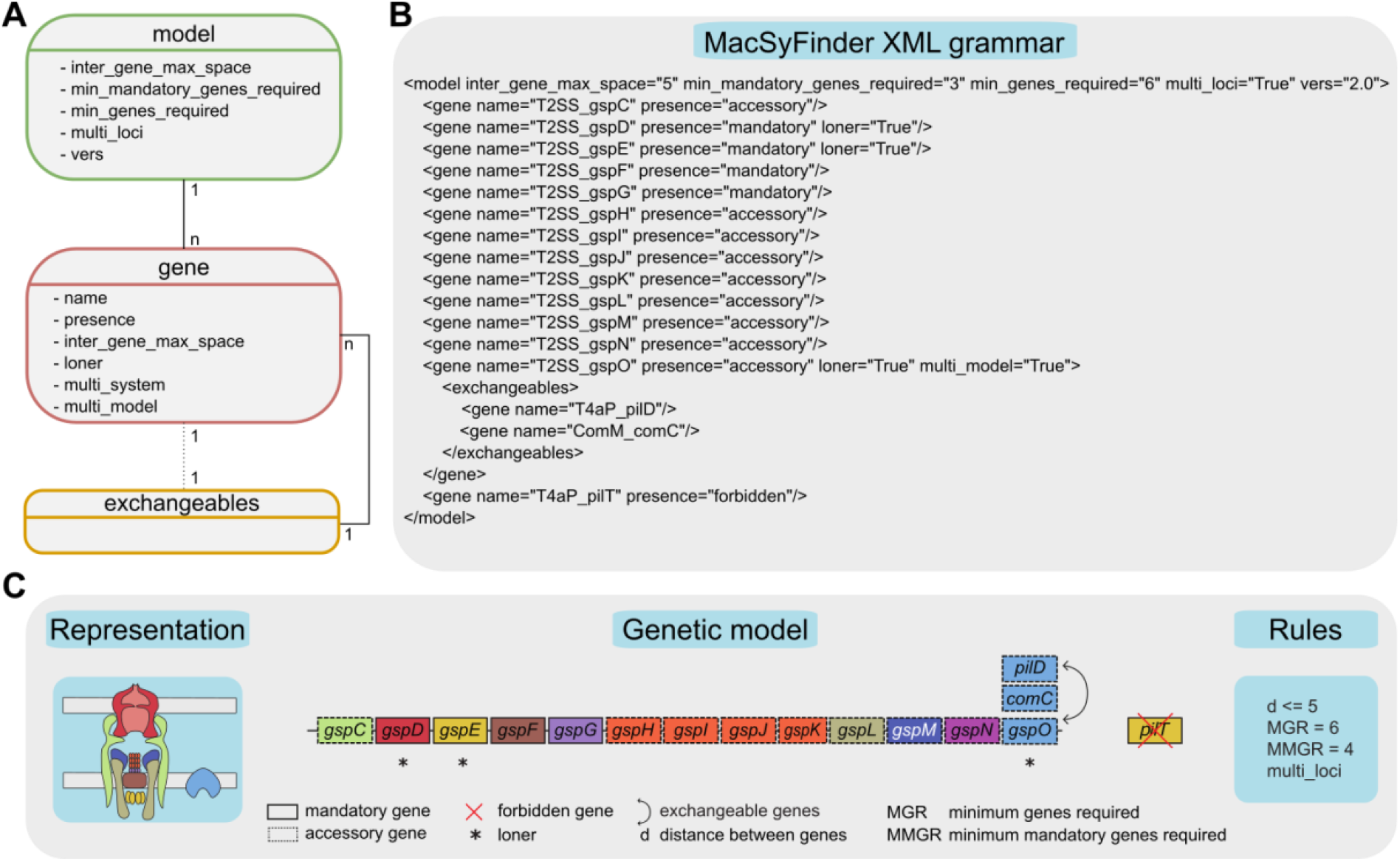
Description of the hierarchical grammar used in MacSyFinder models and example of the T2SS model. **A**. The “model” is the root element of the XML document according to the grammar. It represents the system to model and contains at least one element “gene”. The “gene” element describes the genes constituting a system. It may contain one element “exchangeables”. The dashed line between “gene” and “exchangeables” illustrate the fact that a gene does not necessary contain an “exchangeables” element. The “exchangeables” element contains a set of genes (one at least) that can replace functionally the parent “gene” in the system quorum. The one-to-many relationships between the different elements is represented by lines connecting the boxes, with the cardinality of the relationship appearing next to the element. The diverse possible features of each element are represented in the corresponding boxes. **B**. XML grammar of the T2SS from TXSScan v1.1.1 (and TFFscan v1.0.1) (Abby et al., 2016; Denise et al., 2019). **C**. A schematic representation of the T2SS machinery spanning the membranes of a diderm bacterium is displayed on the left. The genetic model corresponding to the T2SS model in panel B is illustrated in the central part, with gene boxes filled with the colour of the corresponding proteins on the T2SS schema. The quorum and co-localization rules to fulfil the T2SS model are described on the right. Genes’ names were abbreviated in the genetic model compared to the names in the XML model. The C panel was derived from (Denise et al., 2019).

### II/ A new system modelling and search engine for a more relevant exploration of possible systems

#### A systematic comparison of MacSyFinder v1 versus v2

We used TXSScan to systematically compare the results of MacSyFinder version 2 with those of version 1 (Fig. 4 and Table S2). We ran both versions on the same set of genomes, and computed the total number of detected systems, and the average system completeness (proportion of the listed system’s genes detected in a system). As anticipated, the results of both versions were very similar for single-loci systems, yet more systems were detected with v2 (8% increase). A fundamental improvement of the novel version is that the systems are searched one by one: the identified genes are filtered by type of system and assembled in clusters if relevant (Fig. 1). The new search engine can thus resolve much better the most complex cases. It also prevents the spurious elimination of relevant candidate systems, *e*.*g*., when a gene from another system is within a cluster of the candidate system, which is the cause for false negatives in v1 (*e*.*g*. for T6SSi in Fig. 4).

The novel search engine explores the space of possible solutions combinatorically (Fig. 1). This results in noticeable improvements for the detection of multi-loci systems. Hence, MacSyFinder v2 annotated around 20% more multi-loci systems than v1 (Fig. 4B, Table S2, example in section IV).

In addition to a higher number of systems annotated with version 2 (10% more), the annotated systems displayed a similar or higher level of completeness (“wholeness” in Fig. 4C). Overall, these results validate the relevance of the new search algorithm and of the explicit scoring system that we chose to favour complete and concise systems. These advantages are illustrated by examples in Sections III-IV.

**Figure 4.**
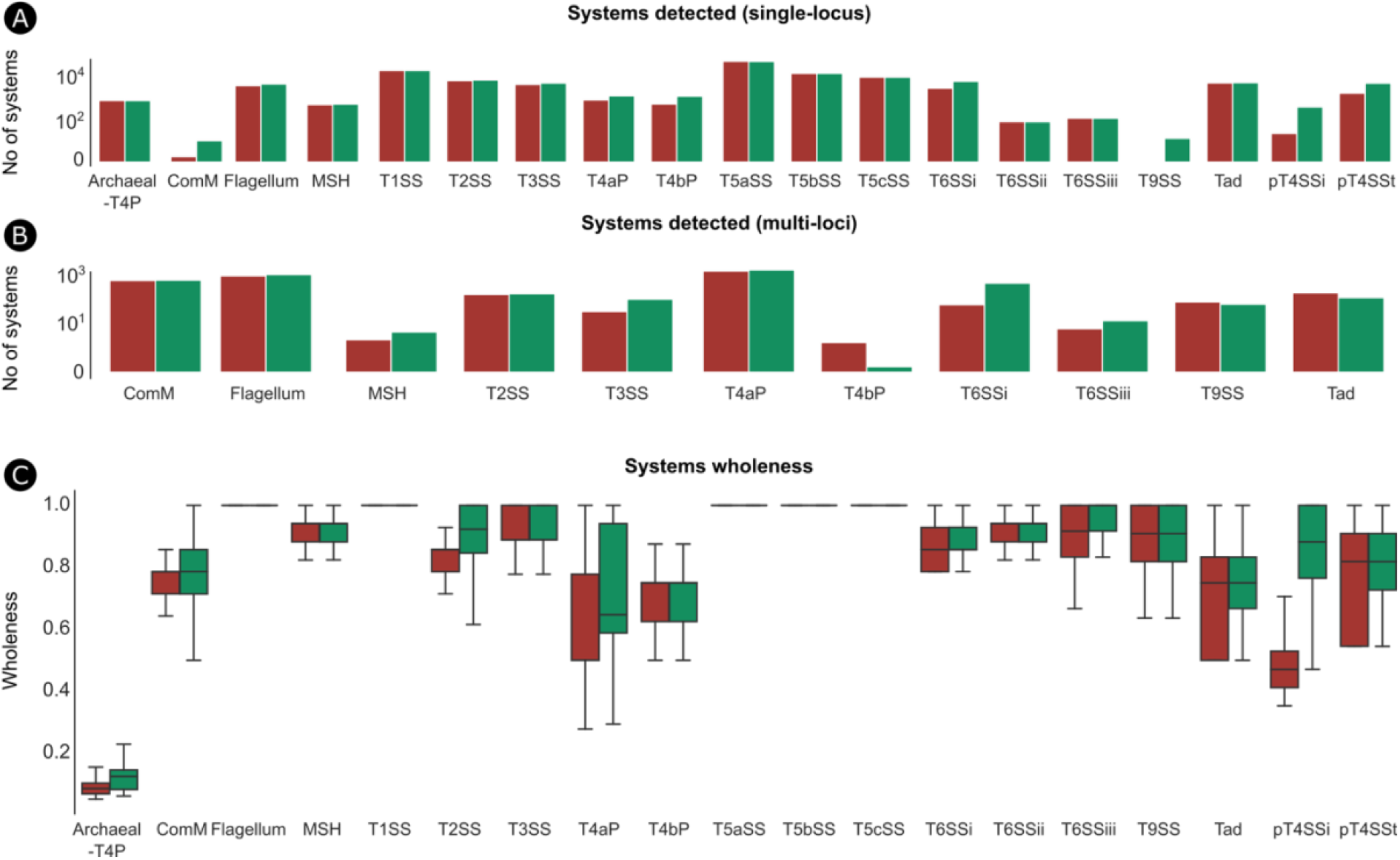
Comparison of MacSyFinder v1 and v2 using TXSScan. MacSyFinder v1 (red) and v2 (green) were used on the same set of complete prokaryotic genomes with the models from TXSScan. Different statistics were used to compare their performance for systems’ detection: **(A)** the number of systems detected as “single-locus” and **(B)** “multi-loci”, and **(C)** a boxplot showing the distribution of the wholeness of detected systems. The wholeness of a detected system is the number of detected genes divided by the number of genes listed as part of the system in the system’s model definition (forbidden and neutral genes excluded).

#### Assessing the performance of MacSyFinder v2

The novel combinatorial examination of sets of genes and clusters to identify candidate systems (see Materials and Methods) can deal with more complex cases, e.g. the occurrence of multiple scattered systems (see section IV for an example). However, the combinatorial exploration may be computationally costly, especially when there are many occurrences of clusters and genes. This cost is partly relieved by filtering the protein hits using the GA scores (or other criteria), because this effectively removes many false positives and leaves fewer genes and clusters to consider. Yet when testing the new search engine, we were sometimes confronted to cases of genomes with dozens of hits for “out-of-cluster” genes (loner or multi_system genes). The analysis of all combinations of such genes can be extremely costly. To make these cases manageable, MacSyFinder v2 uses a heuristic that considers several occurrences of the same “out-of-cluster” gene as a single one representative gene in order to form otherwise equivalent combinations (in quorum and score) for potential systems (Fig. S2). This “representative” is selected as the one with the best matching protein (best HMMER score). The other “out-of-cluster” genes detected are kept and listed separately in dedicated files (best_solution_loners.tsv and best_solution_multisystems.tsv). This makes the combinatorial exploration of solutions manageable in most if not all the cases. If this is not the case, we advise the users to revise their system’s modelling strategy and/or HMM profiles specificity (e.g., increase the GA score thresholds). Finally, even if there is a graph-based search for the best solution (see Materials and Methods and Fig. 2B), MacSyFinder also provides output files with valid systems that are not part of the best solution that may be of interest to the user in some situations (system variants discovery, detection of degraded systems, etc.).

We used the Python profiler (cProfile) to assess MacSyFinder performances and find where the program is more expensive in terms of running time. As anticipated, we found two bottlenecks: the search of genes by HMMER and the resolution of the best solution from combinations of candidate systems. The latter is based on a graph-based search of the maximal clique with NetworkX. This algorithm is known to be exponential. We measured the time spent when running MacSyFinder (with CONJScan, CasFinder and TXSScan, see Materials and Methods) on 6092 bacterial genomes and discarded the genomes where no systems had been found. We analysed 2455, 2241 and 5321 bacterial chromosomes respectively for CONJScan, CasFinder and TXSScan. We could confirm that the computational time grows exponentially with the number of solutions (Fig. S3), or the number of systems’ candidates (Fig. 5A, Fig. S4), and for all three macsy-models used. However, running times to find the best solution were usually small: less than 1 second for all runs with CasFinder and CONJScan models, and for 98% of the runs with TXSScan (Fig. 5B). In some cases, the number of candidates increased tremendously and MacSyFinder could not find a solution within 3 hours. This corresponded to 9 genomes of 6092 with TXSScan. This coincided with the most complicated models (with hundreds of profiles, and multi-loci systems with loners, multi_sytems and multi_model genes). In addition, the 9 genomes were large, encoding between 4053 and 12492 genes. For this reason, we introduced a “--timeout” option (not set by default) to stop the search after a user-defined amount of time. It can be suitable for users who have a collection of genomes to analyse and do not want to be delayed by one, while other users may be interested in obtaining a result even if this means longer run times.

**Figure 5.**
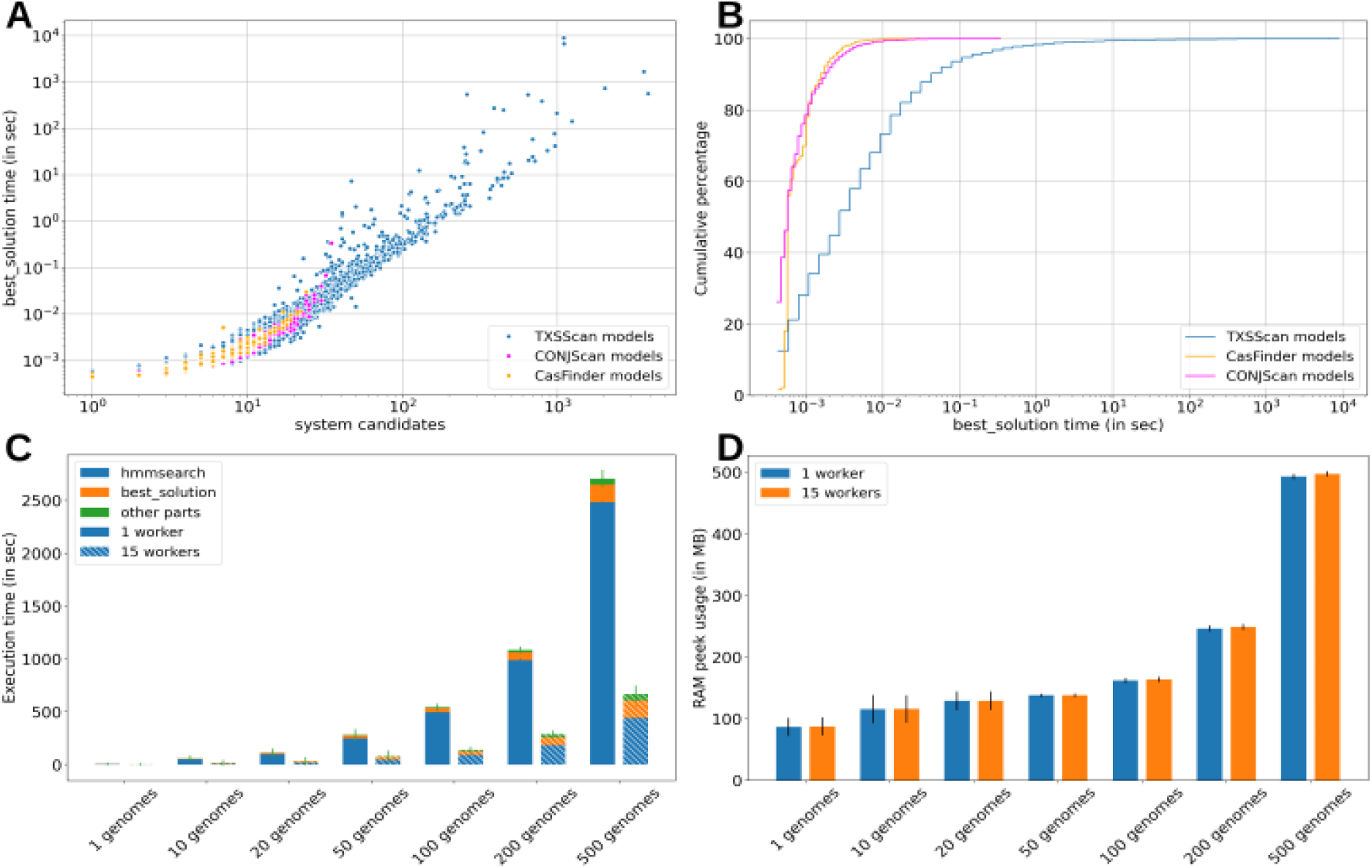
Performance testing of MacSyFinder v2. **A**. The time spent by MacSyfinder in the resolution of the best solution grows exponentially with the number of solutions (Fig. S3) and with the number of candidate systems. **B**. Although the algorithm runs in exponential time, the broad majority of the runs complete the best solution resolution within 1 second. **C**. The mean time spent by MacSyFinder searching genes with hmmsearch (HMMER) is large compared to the other parts of the program, and grows with the number of genomes in the dataset (“gembase” mode). However this time can be significantly reduced when using multiple workers. **D**. The mean maximum resident memory used by MacSyFinder v2 is typically 100MB to analyse one genome and this grows with the size of the dataset to analyse, while being still manageable on a laptop, with less than 500 MB used for 500 genomes analysed. For the 1, 10, 20, 50, 100, 200 and 500 genomes datasets (panels C and D), respectively 1500, 150, 75, 30, 15, 7 and 3 non-redundant replicate datasets were sampled and analysed. Bars in C and D panels report the mean standard error.

The second bottleneck in MacSyFinder is the search of genes with HMMER (hmmsearch) that consists in the most computationally intensive part of a MacSyFinder run (Fig. 5C). The timing of this step grows with the number of genomes in the dataset (“gembase” search mode). To tackle this problem, the user can benefit from available multi-core CPU to search several genes in parallel with the “--worker” option. Using this option, the time spent in the genes search stage decreases significantly, even with a dataset containing a single genome. This technique is very efficient for macsy-models containing numerous genes, such as TXSScan (340 genes) (Fig. 5C).

In addition to the multi-core implementation for the genes search, we designed and made available a parallel version of MacSyFinder that consists in a workflow based on NextFlow (https://www.nextflow.io/) recommended for the analysis of datasets containing many genomes and easily deployed on a computer cluster.

The memory footprint of MacSyFinder v2 for the analysis of one genome is around 100 MB of RAM (with the heavy TXSScan macsy-model), and remains very contained (<500 MB) even when analysing hundreds of genomes (“gembase” mode). This relatively low memory footprint is mainly due to the NetworkX’s maximal clique search algorithm, which is implemented as a generator that never stores all cliques in the memory at once (Fig. 5D & Fig. S5). Overall, the time and RAM used by MacSyFinder makes it easily useable on a laptop even when analysing hundreds of genomes.

We illustrate and discuss in further detail the advantage of the new version of MacSyFinder for the search of molecular systems in the following sections. For this, we applied it to three types of systems: the CRISPR-Cas system (CasFinder package), the conjugative systems (CONJscan package), and the type IV filament super-family (TFFscan and TXSScan packages).

### III/ Application of MacSyFinder v2 to the identification of Cas systems (CasFinder)

CRISPR-Cas systems are adaptive immune systems that protect Bacteria and Archaea from invasive agents (phages, plasmids, etc.) (Hampton et al., 2020). A typical CRISPR-Cas system consists of a CRISPR array and adjacent cluster of *cas* (CRISPR associated) genes that form one or more operons of 1 to 13 genes (Fig. 6A) (Makarova et al., 2020). As CRISPR arrays do not code for proteins, this part of the system is not identified by MacSyFinder. The *cas* genes clusters are very diverse and are currently classified into two classes, six types (I-VI) and more than 30 subtypes based on their composition in *cas* genes (Makarova et al., 2020). We have previously developed a package of models called CasFinder dedicated to the detection of CRISPR-Cas systems using MacSyFinder v1 (Abby et al., 2014; Couvin et al., 2018). We hereby propose an updated and improved version of CasFinder that benefits from the new features of MacSyFinder v2.

**Figure 6.**
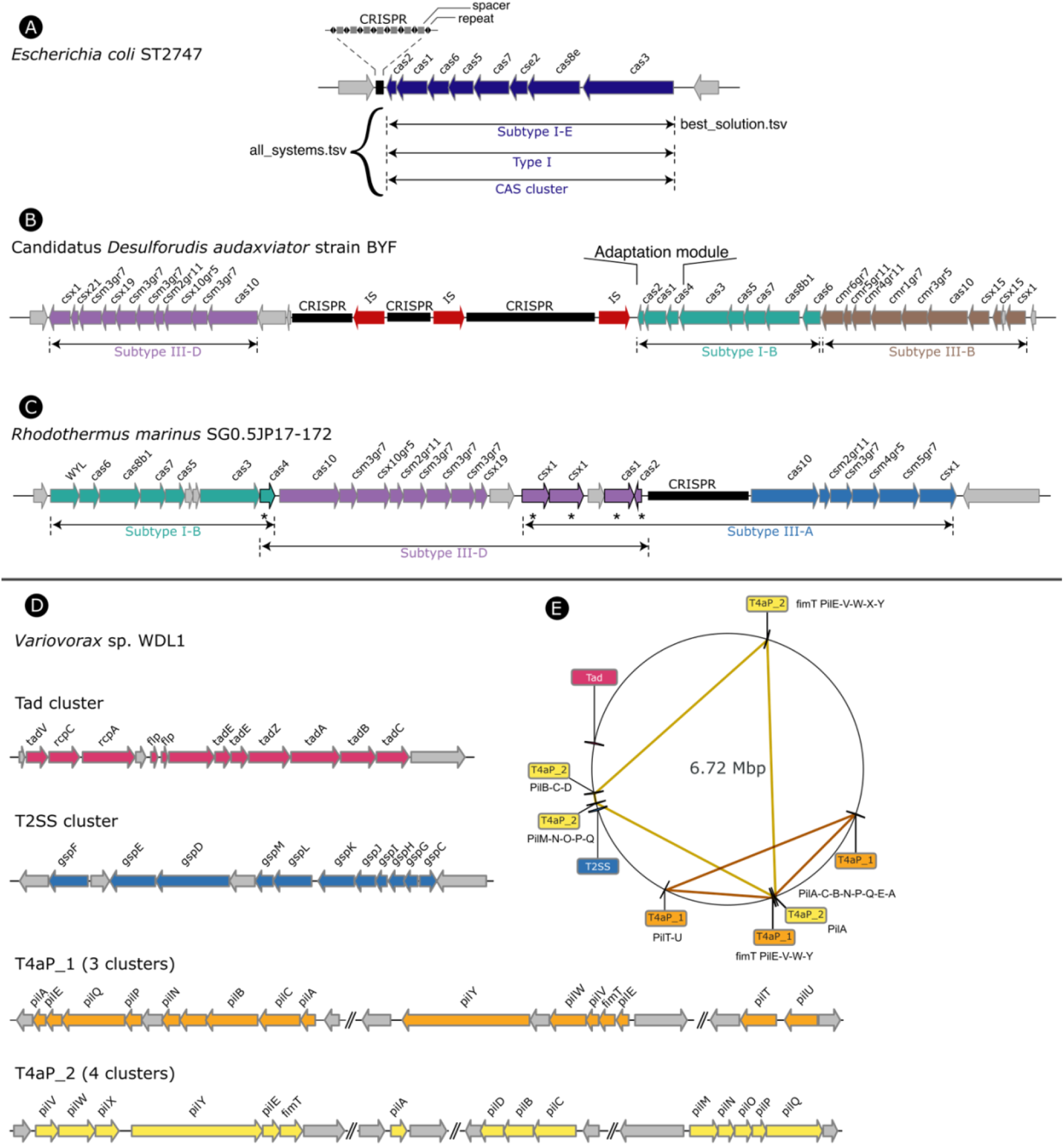
Application of MacSyFinder v2 to CasFinder (v3.1.0) and TFFscan (v1.0.0). **(A)** CRISPR-Cas system has two parts: a CRISPR array and a cluster of *cas* genes. The new MacSyFinder search engine simultaneously annotates Cas clusters at 3 levels of classification from the most accurate (i.e. the subtype level) to the most permissive. When possible, it favours as the best solution the annotation at the subtype level but allows to recover atypical or decayed systems with the 2 other levels of classification. **(B)** The combinatorial approach for the search of the best solution improves the detection of tandem systems. All models are tested and challenged, then the best combination of systems is determined. Here, it reveals the presence of 3 systems of different subtype in tandem (one color per subtype). **(C)** The new search engine avoids overlap between different candidate systems to determine the best solution(s), unless specified in the model with the multi_system or multi_model features. As illustrated, the adaptation module (cas1, cas2 and cas4) has been defined as “multi_model” (indicated by a star*) in some subtype models and can thus be assigned to 2 systems in tandem, which improves their identification. Without this feature newly implemented in v2, one of the two systems would be lost. **(D)** Several members of the Type IV filament super-family (TFF-SF) could be found in the genome of *Variovorax* sp. WDL1. The new search engine enables the annotation of two distinct T4aP in the same genome. Here we can observe that the two detected T4aP are gathering clusters of different and complementary gene composition, underlying their coherence. The two strokes between each gene cluster signifies that the clusters are not close to each other on the chromosome. **(E)** The location of the T4aP gene clusters is displayed along the circular chromosome. A polygon connects the different parts of a same system with colors matching that of the systems in panel D. In all panels, genes are represented by arrows, their length indicates the gene length, and their direction indicates the gene orientation.

#### The graph-based approach improves tandem systems detection

CRISPR-Cas systems can be subdivided into three distinct, though partially overlapping, functional modules. Some of these, e.g., the adaptation module mainly composed of Cas1, Cas2, and Cas4 proteins, may be very similar between subtypes or even types, making the detection of tandem systems particularly challenging. With the v2 new search engine, all systems are searched one by one. The best possible combination of systems is retrieved using a graph-based approach, which significantly improves the identification of tandem systems. This improvement is even more important when the number of tandem systems exceeds two, as MacSyFinder v1 could not handle these rare complex situations at the subtype level (Fig. 6B).

#### The “multi_model” gene feature enables tandem, overlapping systems detection

Most CRISPR-Cas systems have an adaptation module when they are alone. But, when they are in tandem, it is not uncommon to find that only one module is present for both systems (Bernheim et al., 2019). This complicates its detection, especially when it is located between tandem systems. In v1, the adaptation module was assigned to one of the two systems at the risk of missing the second one if the latter turned out to be too small (i.e., with a minimum number of required genes lower than the defined threshold). In v2, thanks to the new “multi_model*”* gene feature, it is possible to allow a gene to be present in different models. Thus, by defining the proteins involved in the adaptation module as “multi_model”, they are assigned to the two overlapping systems (Fig. 6C).

#### The new search engine and scoring system allow searching for different levels of classification simultaneously

Some Cas subtypes are extremely similar in terms of gene content and require very precise decision rules to distinguish them. However, the more precise these rules are, the higher the risk is of missing systems. To overcome this difficulty, we have previously defined different sets of models providing detection at three levels of classification, from the most permissive to the most specific one: (1) a general model (called CAS_cluster) allowing the identification of any cluster of *cas* genes, (2) a set of models for detection at the type level, (3) and finally a set of models for detection at the subtype level. MacSyFinder v1 analyzed the models one by one in a pre-defined order and selected the first model whose rules were satisfied. Using all three sets of models simultaneously meant that not all possibilities were explored. Thanks to the new v2 search engine and scoring system, all models can now be analyzed at once. Therefore, it is possible for the same cluster to be detected at several classification levels (Fig. 6A). The choice of the best solution presented to the user among these different assignments is based on the score of each candidate, then on their wholeness (proportion of genes found over the number of listed ones, or over “max_nb_genes” if defined in the model). Here, the subtype level models have been defined with a “max_nb_genes” parameter lower than for the models higher in the classification. Thus, for a given system that will obtain the same score from several classification levels, the most specific one will obtain the higher system’s wholeness, ensuring the most specific annotation is proposed as the best solution. We thus favoured annotation at the subtype level as being by far the most informative. Still, when the subtype-level search fails, the program allows the detection of atypical or decayed clusters via the models at the other levels (type-level or general case).

### IV/ Application of MacSyFinder v2 to TFFscan and CONJscan

#### The new search engine and scoring system allow the retrieval of various occurrences of Type IV pili encoded at multiple loci

The type IV filaments super-family (TFF-SF) is a family of homologous machineries involved in bacterial and archaeal motility (e. g., the type IVa pilus “T4aP” and archaeal flagellum), toxin secretion (e.g., type II secretion systems, T2SS) or exogenous DNA acquisition (e. g., competence apparatus, Com) (Pelicic, 2008). Some members of the TFF-SF have their genes scattered across the genome (e.g., T4aP and some T2SS), and some genomes may harbour several scattered occurrences of the same system (Denise et al., 2019). In this case, it is not trivial to identify and discriminate the occurrences of the different systems. The search engine of MacSyFinder v1 collected occurrences of the same system as one large system containing multiple copies of several genes. The new v2 search engine examines and then scores all possible combinations of gene clusters and (authorized) out-of-cluster genes eligible as systems. The scoring of these candidate systems penalizes the presence of the same gene in several gene clusters. This approach favours solutions presenting complete yet concise systems. For example, it allows the separation of two different multi-loci T4aP found in the same genome (Fig. 6D-E).

#### The new scoring scheme allows to distinguish putatively decayed conjugative elements from the others

Integrative conjugative elements and conjugative plasmids are very abundant mobile genetic elements that can transfer themselves from one bacterium to another. To do so, they encode a conjugative system that includes a relaxase (MOB) and a mating pair formation (MPF) machinery responsible for pilus biogenesis and mating junctions (de la Cruz et al., 2010). The known relaxases are currently searched using 11 HMM profiles, and the MPFs are classified into eight different types (FA, FATA, B, C, F, G, I, and T). Together, they make for eight models of T4SS (Guglielmini et al., 2013). MPFs include numerous genes, between eight to several dozens. However, the conserved set of genes seen as mandatory is much smaller (relaxase, VirB4, coupling protein), and the other conserved genes oscillate between seven and 27. From wet-lab experiments to pandemic studies or phylogenetic analyses, discriminating between complete transferrable elements and incomplete, potentially immobile conjugative elements, is crucial. We have recently shown that decayed conjugative elements are not rare (Coluzzi et al., 2022). Hence, it would be important to have an easy way to identify complete and incomplete systems. To tackle this problem, we developed macsy-models taking advantage of the new scoring scheme implemented in v2.

All systems can be tested and challenged at once in the new version. The selection of the best solution among different candidate systems is based on the score of each candidate. Using this new feature, we created models designed to compete with each other (Fig. 7). For each conjugative system, one model was designed to detect complete systems, while the other was designed to detect both complete and incomplete systems. Used independently, the complete model would only detect complete systems, and the incomplete model would indiscriminately detect complete and incomplete systems. However, when used together in competition, the scoring system attributes actual complete systems to the complete model while incomplete systems are only detected by the “incomplete” model (Fig. 7).

**Figure 7.**
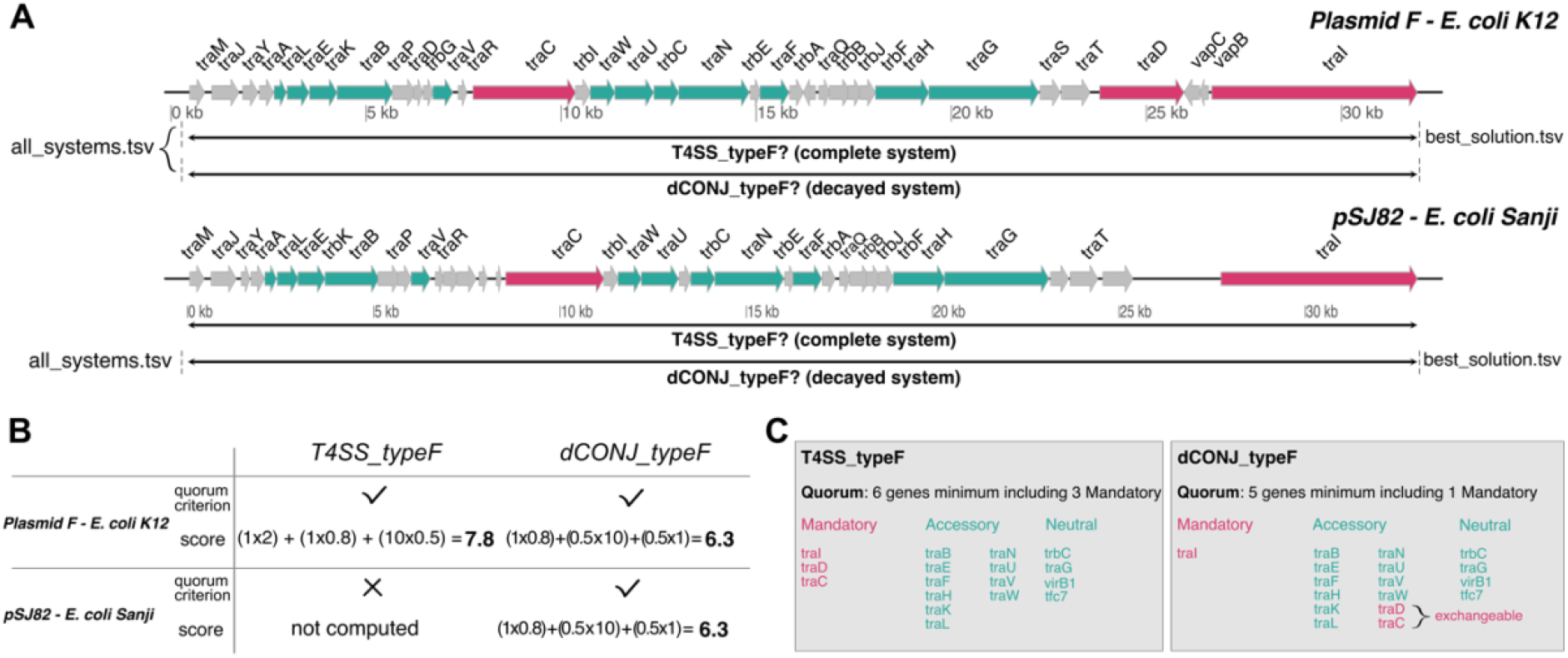
Application of MacSyFinder v2 to distinguish complete and incomplete conjugative systems on bacterial plasmids with CONJscan v2.0.1. **A**. Representation of a complete conjugative system (top) and a decayed conjugative system (bottom). Arrows represent the predicted genes of the plasmids and their orientation. Mandatory and accessory genes of the systems are represented in fuchsia and cyan respectively. **B**. Description of the score for the complete MPF_F_ (T4SS_TypeF) model and “decayed” MPF_F_ model (dCONJ_typeF) when computed by the scoring scheme of MacSyFinder v2 (detailed in Fig. 2A). CONJScan plasmids’ models were used all at once with the “all” option. **C**. Difference between the complete and decayed models for the MPF_F_. Both models list the same genes, but the required quorum of mandatory genes and total genes required are different. The model designed to detect complete systems (T4SS_typeF) requires 3 mandatory genes and 6 genes minimum, while the “decayed” model (dCONJ_typeF) was designed to require only 1 mandatory gene and the other mandatory genes were set as “accessory” and exchangeable between each other. Thus, we ensure that if the quorum is reached for a complete system, the score of the “complete” model is always higher than the score of the “decayed” model.

## CONCLUSION

MacSyFinder leverages the power of comparative genomics for accurate system-level annotation of microbial genomes. MacSyFinder version 2 enables more relevant and comprehensive system modelling and search capacities. The variety of the applications illustrated here and elsewhere demonstrates the potential of MacSyFinder to annotate many other cellular functions, including biosynthetic gene clusters, metabolic and signalling pathways. The *macsydata* tool and “MacSy Models” Github organization allow systems’ modellers to easily share their macsy-model packages. We hope this will increase the visibility of their contribution and enhance the development of novel models for other molecular systems.

## DATA, SCRIPT AND CODE AVAILABILITY

MacSyFinder source code and the hereby presented macsy-model packages are available at the following Github repositories: https://github.com/gem-pasteur/macsyfinder and https://github.com/macsy-models. A snapshot of MacSyFinder, release v2.1 as described in this article is available at the Software Heritage Archive at the following permalink: https://archive.softwareheritage.org/swh:1:snp:be6f94d82f11702214af35de13a426a280c77353;origin=https://github.com/gem-pasteur/macsyfinder.

Two sets of examples with corresponding command lines and expected input and output files are provided on the Figshare platform: https://doi.org/10.6084/m9.figshare.21581280 and https://doi.org/10.6084/m9.figshare.21716426.v1.

## SUPPLEMENTARY INFORMATION

All supplementary data, code and information (tables and figures) are available from the following Figshare repository: https://doi.org/10.6084/m9.figshare.21936992. The file **macsyfinder_2.1.zip** contains an archive of the described MacSyFinder v2.1 source code. The file **MSF2_Supplementary.pdf** contains the supplementary tables and figures listed in main text. The file **DatasetS1.tsv** contains the list of genomes analysed in this article.

## AUTHORS’ CONTRIBUTIONS

BN, EPCR and SSA designed the new version of MacSyFinder. BN conceived the software architecture and design, the test design, performed the implementation and the performance tests. RD, CC, MT, EPCR and SSA tested MacSyFinder. RD, CC, MT and SSA updated and distributed the presented macsy-model packages on the dedicated repository. RD, CC and MT analysed the results of MacSyFinder detection and implemented GA scores within HMM profiles of the presented macsy-model packages. EPCR and SSA wrote the first versions of the manuscript, and all authors contributed to and approved the final versions of the manuscript.

## ACKOWLEDGEMENTS

The authors are grateful to Yoann Dufresne for the suggestion to use the NetworkX library to address the weighted maximum clique search problem. The authors thank Amandine Perrin for providing the genomes’ lists for the performance tests. This work used the computational and storage services (TARS cluster) provided by the IT department at Institut Pasteur, Paris.

## REFERENCES

Abby SS, Cury J, Guglielmini J, Néron B, Touchon M, Rocha EPC (2016) Identification of protein secretion systems in bacterial genomes. Scientific Reports, 6, 23080. https://doi.org/10.1038/srep23080

Abby SS, Denise R, Rocha EP (2023) Identification of protein secretion systems in bacterial genomes using MacSyFinder version 2. BioRxiv preprint. https://doi.org/10.1101/2023.01.06.522999

Abby SS, Neron B, Menager H, Touchon M, Rocha EP (2014) MacSyFinder: a program to mine genomes for molecular systems with an application to CRISPR-Cas systems. PLoS One, 9, e110726. https://doi.org/10.1371/journal.pone.0110726

Abby SS, Rocha EPC (2012) The Non-Flagellar Type III Secretion System Evolved from the Bacterial Flagellum and Diversified into Host-Cell Adapted Systems. PLoS Genet, 8, e1002983. https://doi.org/10.1371/journal.pgen.1002983

Adam PS, Borrel G, Gribaldo S (2019) An archaeal origin of the Wood-Ljungdahl H4MPT branch and the emergence of bacterial methylotrophy. Nat Microbiol, 4, 2155–2163. https://doi.org/10.1038/s41564-019-0534-2

Bernheim A, Bikard D, Touchon M, Rocha EPC (2019) Atypical organizations and epistatic interactions of CRISPRs and cas clusters in genomes and their mobile genetic elements. Nucleic Acids Research, gkz1091. https://doi.org/10.1093/nar/gkz1091

Blin K, Shaw S, Kloosterman AM, Charlop-Powers Z, van Wezel GP, Medema MH, Weber T (2021) antiSMASH 6.0: improving cluster detection and comparison capabilities. Nucleic Acids Research, 49, W29–W35. https://doi.org/10.1093/nar/gkab335

Brandes U, Erlebach T (2005) Network Analysis. Methodological Foundations. (U Brandes, T Erlebach, Eds,). Springer Berlin, Heidelberg.

Chibani CM, Mahnert A, Borrel G, Almeida A, Werner A, Brugere JF, Gribaldo S, Finn RD, Schmitz RA, Moissl-Eichinger C (2022) A catalogue of 1,167 genomes from the human gut archaeome. Nat Microbiol, 7, 48–61. https://doi.org/10.1038/s41564-021-01020-9

Coluzzi C, Garcillan-Barcia MP, de la Cruz F, Rocha EPC (2022) Evolution of Plasmid Mobility: Origin and Fate of Conjugative and Nonconjugative Plasmids. Mol Biol Evol, 39. https://doi.org/10.1093/molbev/msac115

Couvin D, Bernheim A, Toffano-Nioche C, Touchon M, Michalik J, Neron B, Rocha EPC, Vergnaud G, Gautheret D, Pourcel C (2018) CRISPRCasFinder, an update of CRISRFinder, includes a portable version, enhanced performance and integrates search for Cas proteins. Nucleic Acids Res, 46, W246–W251. https://doi.org/10.1093/nar/gky425

de la Cruz F, Frost LS, Meyer RJ, Zechner EL (2010) Conjugative DNA metabolism in Gram-negative bacteria. FEMS Microbiol Rev, 34, 18–40. https://doi.org/10.1111/j.1574-6976.2009.00195.x

Cury J, Abby SS, Doppelt-Azeroual O, Néron B, Rocha EPC (2020) Identifying Conjugative Plasmids and Integrative Conjugative Elements with CONJscan. In: Horizontal Gene Transfer (ed de la Cruz F), pp. 265–283. Springer US, New York, NY. https://doi.org/10.1007/978-1-4939-9877-7_19

Cury J, Touchon M, Rocha EPC (2017) Integrative and conjugative elements and their hosts: composition, distribution and organization. Nucleic Acids Research, 45, 8943–8956. https://doi.org/10.1093/nar/gkx607

Dandekar T, Snel B, Huynen M, Bork P (1998) Conservation of gene order: a fingerprint of proteins that physically interact. Trends in Biochemical Sciences, 23, 324–328. https://doi.org/10.1016/s0968-0004(98)01274-2

Denise R, Abby SS, Rocha EPC (2019) Diversification of the type IV filament superfamily into machines for adhesion, protein secretion, DNA uptake, and motility. PLOS Biology, 17, e3000390. https://doi.org/10.1371/journal.pbio.3000390

Denise R, Abby SS, Rocha EPC (2020) The Evolution of Protein Secretion Systems by Co-option and Tinkering of Cellular Machineries. Trends in Microbiology, 28, 372–386. https://doi.org/10.1016/j.tim.2020.01.005

Eddy SR (2011) Accelerated Profile HMM Searches. PLoS Comput Biol, 7, e1002195. https://doi.org/10.1371/journal.pcbi.1002195

Guglielmini J, de la Cruz F, Rocha EPC (2013) Evolution of conjugation and type IV secretion systems. Molecular biology and evolution, 30, 315–331. https://doi.org/10.1093/molbev/mss221

Haft DH, Selengut JD, White O (2003) The TIGRFAMs database of protein families. Nucleic Acids Research, 31, 371–373. https://doi.org/10.1093/nar/gkg128

Hagberg AA, Schult DA, Swart PJ (2008) Exploring Network Structure, Dynamics, and Function using NetworkX. In: (eds Varoquaux G, Vaught T, Millman J)

Hampton HG, Watson BNJ, Fineran PC (2020) The arms race between bacteria and their phage foes. Nature, 577, 327–336. https://doi.org/10.1038/s41586-019-1894-8

Huynen M, Snel B, Lathe W, Bork P (2000) Predicting protein function by genomic context: quantitative evaluation and qualitative inferences. Genome Res, 10, 1204–10. https://doi.org/10.1101/gr.10.8.1204

Kanehisa M, Sato Y (2019) KEGG Mapper for inferring cellular functions from protein sequences. Protein Sci. https://doi.org/10.1002/pro.3711

Karp PD, Paley SM, Midford PE, Krummenacker M, Billington R, Kothari A, Ong WK, Subhraveti P, Keseler IM, Caspi R (2020) Pathway Tools version 24.0: Integrated Software for Pathway/Genome Informatics and Systems Biology.

Makarova KS, Wolf YI, Iranzo J, Shmakov SA, Alkhnbashi OS, Brouns SJJ, Charpentier E, Cheng D, Haft DH, Horvath P, Moineau S, Mojica FJM, Scott D, Shah SA, Siksnys V, Terns MP, Venclovas Č, White MF, Yakunin AF, Yan W, Zhang F, Garrett RA, Backofen R, van der Oost J, Barrangou R, Koonin EV (2020) Evolutionary classification of CRISPR–Cas systems: a burst of class 2 and derived variants. Nature Reviews Microbiology, 18, 67–83. https://doi.org/10.1038/s41579-019-0299-x

Pelicic V (2008) Type IV pili: e pluribus unum? Molecular Microbiology, 68, 827–837. https://doi.org/10.1111/j.1365-2958.2008.06197.x

Pende N, Sogues A, Megrian D, Sartori-Rupp A, England P, Palabikyan H, Rittmann SKR, Grana M, Wehenkel AM, Alzari PM, Gribaldo S (2021) SepF is the FtsZ anchor in archaea, with features of an ancestral cell division system. Nat Commun, 12, 3214. https://doi.org/10.1038/s41467-021-23099-8

Perrin A, Rocha EPC (2021) PanACoTA: a modular tool for massive microbial comparative genomics. NAR genomics and bioinformatics, 3, lqaa106. https://doi.org/10.1093/nargab/lqaa106

Rendueles O, Garcia-Garcera M, Neron B, Touchon M, Rocha EPC (2017) Abundance and co-occurrence of extracellular capsules increase environmental breadth: Implications for the emergence of pathogens. PLoS Pathog, 13, e1006525. https://doi.org/10.1371/journal.ppat.1006525

Sharp C, Foster KR (2022) Host control and the evolution of cooperation in host microbiomes. Nat Commun, 13, 3567. https://doi.org/10.1038/s41467-022-30971-8

Sonnhammer EL, Eddy SR, Durbin R (1997) Pfam: a comprehensive database of protein domain families based on seed alignments. Proteins, 28, 405–420. https://doi.org/10.1002/(sici)1097-0134(199707)28:3<405::aid-prot10>3.0.co;2-l

Taib N, Megrian D, Witwinowski J, Adam P, Poppleton D, Borrel G, Beloin C, Gribaldo S (2020) Genome-wide analysis of the Firmicutes illuminates the diderm/monoderm transition. Nat Ecol Evol, 4, 1661–1672. https://doi.org/10.1038/s41559-020-01299-7

Teichmann SA, Babu MM (2002) Conservation of gene co-regulation in prokaryotes and eukaryotes. Trends Biotechnol, 20, 407–10; discussion 410. https://doi.org/10.1016/s0167-7799(02)02032-2

Tesson F, Herve A, Mordret E, Touchon M, d’Humieres C, Cury J, Bernheim A (2022) Systematic and quantitative view of the antiviral arsenal of prokaryotes. Nat Commun, 13, 2561. https://doi.org/10.1038/s41467-022-30269-9

Vallenet D, Calteau A, Dubois M, Amours P, Bazin A, Beuvin M, Burlot L, Bussell X, Fouteau S, Gautreau G, Lajus A, Langlois J, Planel R, Roche D, Rollin J, Rouy Z, Sabatet V, Medigue C (2020) MicroScope: an integrated platform for the annotation and exploration of microbial gene functions through genomic, pangenomic and metabolic comparative analysis. Nucleic Acids Res, 48, D579–D589. https://doi.org/10.1093/nar/gkz926

