## Supplementary Information for "MacSyFinder v2: Improved modelling and search engine to identify molecular systems in genomes"

### Supplementary figures

A macsy-model package adopts the following structure:

```
family_name
   |________metadata.yml
   |________LICENSE
   |________README.md
   |________model_conf.xml
   |________definitions
   |              |________model_1.xml
   |              |________model_2.xml
   |              :
   |
   |________profiles
                  |________geneA.hmm
                  |________geneB.hmm
                  |________geneC.hmm
                  :
```

Example of the TXSScan v1.1 package that includes sub-families:

```
TXSScan
   |________metadata.yml
   |________LICENSE
   |________README.md
   |________model_conf.xml
   |________definitions
   |              |________archaea
   |              |              |________Archaeal-T4P.xml
   |              |
   |              |________bacteria
   |                             |________monoderm
   |                             |              |________ComM.xml
   |                             |
   |                             |________diderm
   |                                            |________T1SS.xml
   |                                            |________T2SS.xml
   |                                            :
   |
   |________profiles
                  |________ComM_comB.hmm
                  |________ComM_comC.hmm
                  :
                  |________T1SS_abc.hmm
                  |________T1SS_mfp.hmm
                  |________T1SS_omf.hmm
                  |________T2SS_gspA.hmm
                  :
```

**<u>Supplementary Figure 1.</u>** The macsy-model package file architecture and the example of the TXSScan v1.1.1 package.

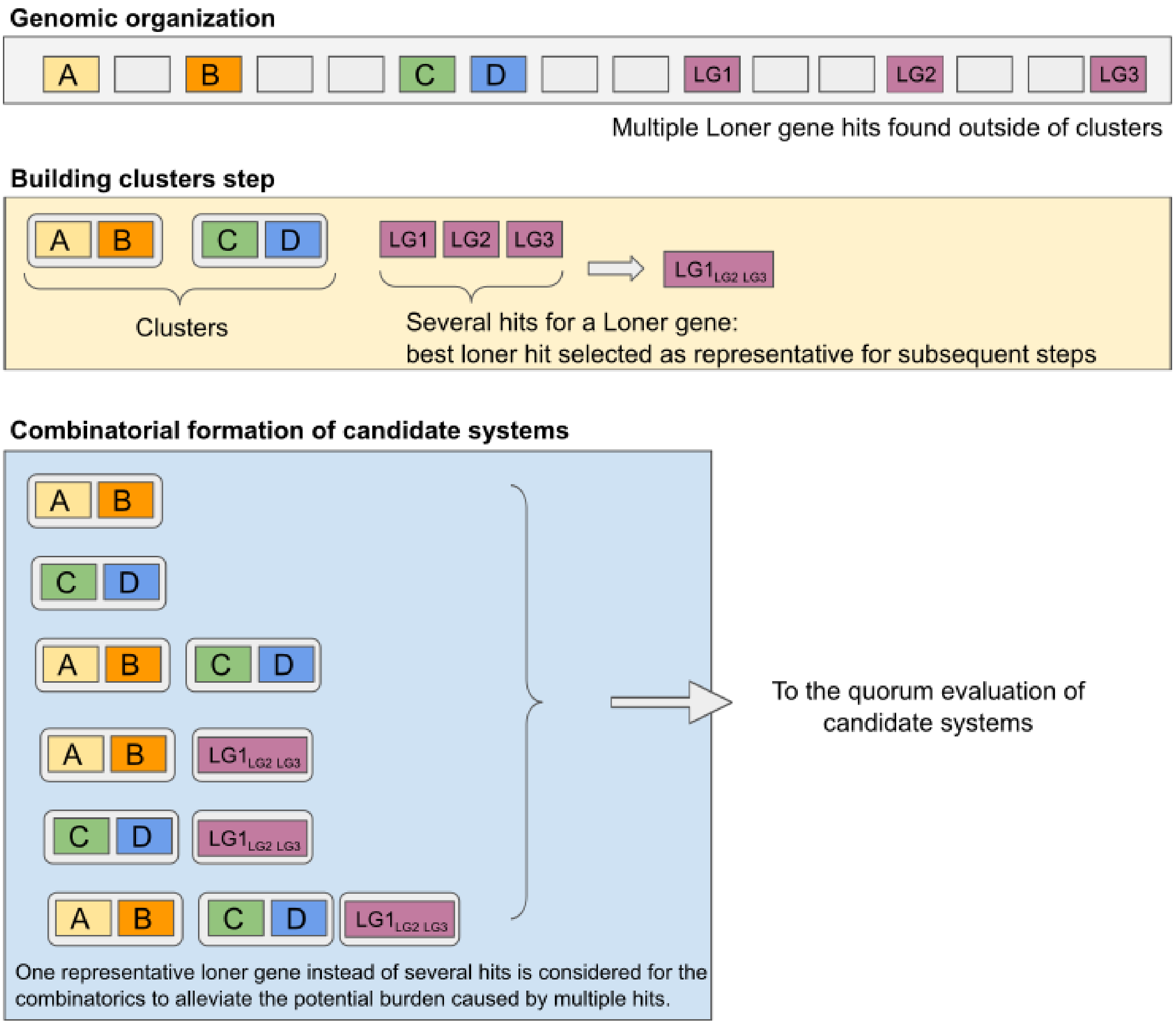


**Supplementary Figure 2.**
MacSyFinder heuristics employed in order to alleviate the burden in combinatorics exploration of multiple out-of-cluster genes (e.g. loner genes “LG”). In a nutshell, in case of multiple out-of-cluster genes, the gene corresponding to the best hit is selected as a representative for the others, and used in place of all of the hits to form candidate systems in combinations with other detected genes and clusters. This enables to reduce the number of evaluated candidate systems, that would anyway be totally equivalent.

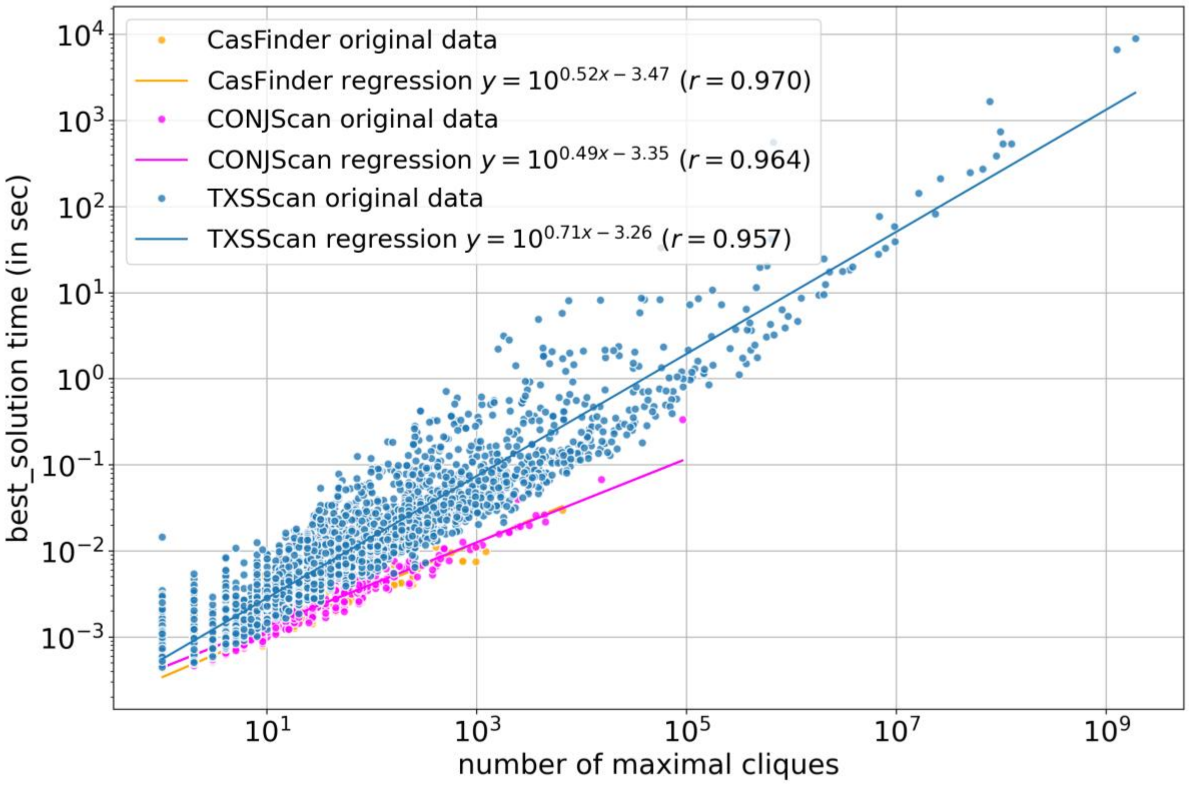


**Supplementary Figure 3.**

The time spent by MacSyFinder in the resolution of the best solution grows exponentially with the number of solutions (maximal cliques). We obtained similar results with three macsy-models tested (CasFinder, CONJScan/chromosome and TXSScan/bacteria). The linear regressions were computed with the “linregress” function from the “stats” module of the “scipy” package. The linear regression was determined from the $log_{10}$ of the number of cliques, and the best solution time.

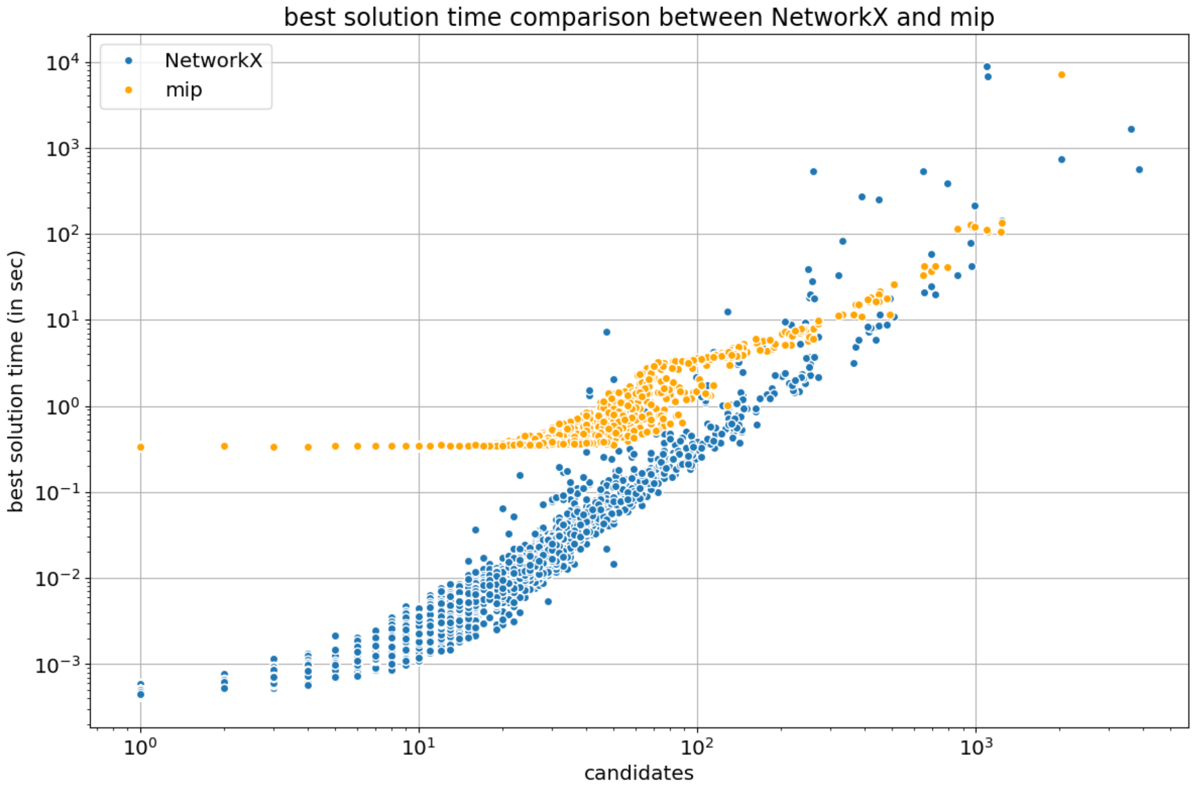


**Supplementary Figure 4.**
We compared the time spent by MacSyFinder to find a best solution (set of compatible systems maximizing the score) with TXSScan on 6453 genomes with two approaches: 1) the *NetworkX* “find_cliques” algorithm where candidate systems define the nodes of an undirected graph and edges connect compatible systems (see Materials and Methods), and 2) by using a Mixed-Integer Linear solver (*Python-MIP* https://www.python-mip.com/). We see that for genomes with high numbers of candidates, both approaches have similar performance in time. But when the number of candidates is low, *NetworkX* is faster than *Python-MIP*. Furthermore, *Python-MIP* gives a single best solution whereas the implementation based on *NetworkX* allows to retrieve all equivalent solutions.

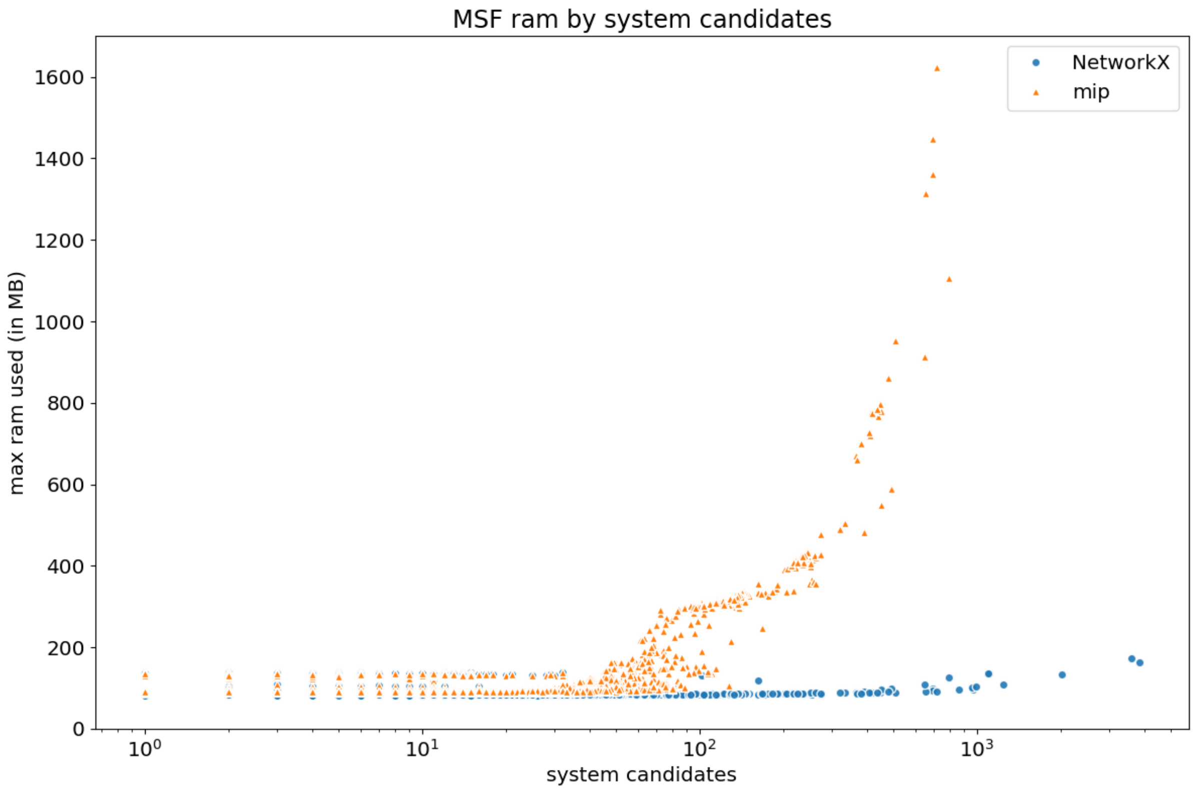


**Supplementary Figure 5.**

We compared the maximum resident memory of a MacSyFinder v2 run when the best solution is solved with either *NetwokX* or *Python-MIP* using TXSScan on 6453 genomes. The *NetworkX* implementation keeps the RAM usage low even when the number of candidates grows. This was not the case with *Python-MIP* where the memory grew exponentially with the number of candidates reaching very high memory usages, eventually resulting in one run being killed by the operating system as it claimed too much memory (> 64GB).

### Supplementary Table

**<u>Supplementary Table 1.</u>** Allowed attributes in the hierarchy of MacSyFinder v2 XML grammar for systems' modelling. Mandatory attributes are in bold

| Element | Attribute | Description | Value |
|---|---|---|---|
| model | **inter_gene_max_space** | An integer representing the maximal number of genes without a match between two genes with a match for a gene's protein profile in order to consider them contiguous and part of a same Cluster of genes | Integer |
| | min_mandatory_genes | An integer representing the minimal number of mandatory genes required to infer the system's presence | Default value to the number of listed mandatory Genes |
| | min_genes_required | An integer representing the minimal number of mandatory or accessory genes required to infer the system's presence | Default value to the number of listed mandatory Genes |
| | multi_loci | If true this allows the definition of "scattered" systems (i.e., systems encoded at different genomic loci or by different gene clusters) | "1", "true", or "0", "false" Default to false |
| | **vers** | The version of the XML grammar | "2.0" |
| | max_nb_genes | An integer representing the maximal number of genes required to consider the system as complete. It is not used for the quorum rules, but it is used as the denominator when calculating the system wholeness (number of genes over number of listed genes, or over max_nb_genes if defined) | Default value to the number of listed Genes (mandatory and accessory) |
| gene | **name** | Name of the gene which must match that of a protein profile enclosed in the "profile" directory of the macsy-model package | String |
| | **presence** | Status of the gene's presence in the system | "mandatory", "accessory", "neutral" or "forbidden" |
| | loner | A *loner* gene can be isolated on the genome and does not have to be part of a cluster of genes to be considered for system's assessment | "1", "true", or "0", "false" Default to false |
| | multi_system | A *multi_system* gene can participate to several occurrences of the current system in the genome | "1", "true", or "0", "false" Default to false |
| | multi_model | A *multi_model* gene can participate to systems from different models. It enables systems presenting this gene to be considered "compatible" at the step of the graph-search for the best solution | "1", "true", or "0", "false" Default to false |

| | inter_gene_max_space | An integer that defines a gene-wise value for "inter_gene_max_space". It supersedes the system-wise parameter | Integer |
|---|---|---|---|
| | exchangeables | Keyword used at the gene-level to list genes that can be used to fulfil the role of the gene of higher level | |

#### Supplementary Table 2. Comparison between MacSyFinder v1 and v2: number of systems detected with TXSScan.

| Systems | Locus types | V1 | V2 |
|---|---|---|---|
| **Archaeal-T4P** | **multi_loci** | 0 | 0 |
| | **single_locus** | 987 | 979 |
| **ComM** | **multi_loci** | 3648 | 3756 |
| | **single_locus** | 4 | 19 |
| **Flagellum** | **multi_loci** | 5750 | 6448 |
| | **single_locus** | 4230 | 4961 |
| **MSH** | **multi_loci** | 13 | 27 |
| | **single_locus** | 657 | 687 |
| **T1SS** | **multi_loci** | 0 | 0 |
| | **single_locus** | 18651 | 18586 |
| **T2SS** | **multi_loci** | 967 | 1042 |
| | **single_locus** | 6876 | 7294 |
| **T3SS** | **multi_loci** | 192 | 620 |
| | **single_locus** | 4746 | 5425 |
| **T4aP** | **multi_loci** | 9005 | 10087 |
| | **single_locus** | 1037 | 1550 |
| **T4bP** | **multi_loci** | 10 | 1 |
| | **single_locus** | 693 | 1506 |
| **T5aSS** | **multi_loci** | 0 | 0 |
| | **single_locus** | 45862 | 45282 |
| **T5bSS** | **multi_loci** | 0 | 0 |
| | **single_locus** | 14066 | 14092 |
| **T5cSS** | **multi_loci** | 0 | 0 |
| | **single_locus** | 9705 | 9667 |
| **T6SSi** | **multi_loci** | 365 | 2835 |
| | **single_locus** | 3295 | 6441 |
| **T6SSii** | **multi_loci** | 0 | 0 |
| | **single_locus** | 123 | 122 |
| **T6SSiii** | **multi_loci** | 37 | 80 |
| | **single_locus** | 173 | 172 |
| **T9SS** | **multi_loci** | 476 | 385 |
| | **single_locus** | 0 | 24 |

| | | | |
|---|---|---|---|
| **Tad** | **multi_loci** | 1123 | 712 |
| | **single_locus** | 5491 | 5656 |
| **pT4SSi** | **multi_loci** | 0 | 0 |
| | **single_locus** | 39 | 517 |
| **pT4SSt** | **multi_loci** | 0 | 0 |
| | **single_locus** | 2017 | 5368 |

#### Supplementary Dataset

**Dataset S1.** List of the replicons analysed with full names and assembly accession identifiers at the NCBI (complete Bacterial and Archaeal genomes, Refseq March 2021). The column “in_test_perf” indicates the sub-set of genomes used for the performance analyses. It is available here: https://doi.org/10.6084/m9.figshare.21936992
